# Inhibitory control of locomotor statistics in walking *Drosophila*

**DOI:** 10.1101/2024.04.15.589655

**Authors:** Hannah C. Gattuso, Karin A. van Hassel, Jacob D. Freed, Kavin M. Nuñez, Beatriz de la Rea, Christina E. May, G. Bard Ermentrout, Jonathan D. Victor, Katherine I. Nagel

## Abstract

In order to forage for food, many animals regulate not only specific limb movements but the statistics of locomotor behavior over time, switching between long-range dispersal and localized search depending on resource availability. How pre-motor circuits regulate such locomotor statistics is not clear. Here we analyze and model locomotor statistics in walking *Drosophila*, and their modulation by attractive food odor. Odor evokes three motor regimes in flies: baseline walking, upwind running during odor, and search behavior following odor loss. During search behavior, we find that flies adopt higher angular velocities and slower ground speeds, and tend to turn for longer periods of time in one direction. We further find that flies spontaneously adopt periods of different mean ground speed, and that these changes in state influence the length of odor-evoked runs. We next developed a simple model of neural locomotor control that suggests that contralateral inhibition plays a key role in regulating the statistical features of locomotion. As the fly connectome predicts decussating inhibitory neurons in the lateral accessory lobe (LAL), a pre-motor structure, we gained genetic access to a subset of these neurons and tested their effects on behavior. We identified one population of neurons whose activation induces all three signature of search and that bi-directionally regulates angular velocity at odor offset. We identified a second group of neurons, including a single LAL neuron pair, that bi-directionally regulate ground speed. Together, our work develops a biologically plausible computational architecture that captures the statistical features of fly locomotion across behavioral states and identifies potential neural substrates of these computations.

## Introduction

To search for food, animals must control both the instantaneous movement of their limbs as well as statistical features of locomotion that alter their overall path through their environment. For example, in seabirds, search trajectories become increasingly localized in response to prey encounters (Paiva et al., 2010; Sommerfeld et al., 2013). Beetles and hoverflies exhibit similar strategies of area-restricted search (Banks, 1957; Chandler, 1969; Fleschner, 1950; Murdie and Hassell, 1973; Dorfman et al., 2022). The nematode *c. elegans* shifts between wide-spread exploratory runs and localized dwelling in response to changes in the availability of food (Ben Arous et al., 2009; Shtonda and Avery, 2006). Similarly, both larval and adult *Drosophila* alter turn statistics in response to changes in odor concentration (Gomez-Marin et al. 2011, Davies et al 2015, Steck et al 2013, Alvarez-Salvado et al 2018, Demir et al. 2020, Stupski et al. 2023). How motor and premotor circuitry are organized to regulate these statistical features of locomotion is not clear.

In recent years, walking adult *Drosophila* has emerged as a powerful model for investigating the neural control of locomotion. In fruit flies, roughly 350 to 500 pairs of descending neurons (DNs) carry motor signals from the brain to the ventral nerve cord, the insect equivalent of the spinal cord (Hsu and Bhandawat, 2016, Namiki et al, 2018, Cheong et al. 2024). Individual DNs have been identified that generate forward walking, turning, or both, when activated (Cande et al. 2018, Rayshubskiy et al. 2024, Bidaye et al. 2020, Yang et al. 2024, Feng et al. 2024). Other DNs promote stopping behavior (Cande et al. 2018, Carreira-Rosasrio et al. 2018, Lee and Doe, 2021, Sapkal et al. 2024). While a small handful of DNs appear to function as “command-like neurons” and generate specific locomotor gestures, imaging studies suggest that many more DNs participate in locomotor control (Yang et al. 2024, Aymanns et al. 2022, Bresovec et al. 2023, 2024, Feng et al. 2024). Further, graded activity in specific DNs or brain regions often correlates with locomotor features; in particular, bilateral activity often correlates with forward velocity while differences in activity across the two brain hemispheres correlate with angular velocity to the left or right (Bidaye et al. 2020, Rayshubskiy et al. 2024, Yang et al. 2024, Bresovec et al. 2023, 2024). Finally, activation of command-like DNs in headless flies argues that interactions between DNs within the brain are crucial to generate coordinated behaviors such as walking and turning (Braun et al. 2024). Together these studies argue that locomotor features such as forward and angular velocity must be encoded by population activity across many DNs.

Complementing this experimental dissection of locomotion, many recent studies have sought to build computational models of walking behavior in flies. Several studies have used Hidden Markov Models to identify behavioral motifs during walking (Berman et al 2016, Katsov et al 2017, Tao et al 2019, Tao et al 2020, Calhoun et al 2019). These models assume that walking behavior can be parsed into discrete motifs that evolve at a set of specific timescales. In contrast, dimensionality reduction approaches suggest that fly walking evolves in a continuous state space (DeAngelis et al. 2019, York et al. 2022), while studies of odor-evoked navigation argue that walking behavior can be modulated at multiple timescales by sensory input (Alvarez-Salvado et al. 2018, Demir et al. 2020, Jayaram et al. 2023). Computational models of walking that feature biologically plausible architectures and generate continuous but low-dimensional variation in walking behavior have been lacking.

In this work, we leverage odor driven behavior to investigate and model the neural control of locomotor statistics. We find that search behavior driven by odor loss involves changes in both ground speed and angular velocity distributions, as well as an increase in unidirectional turning that alters the correlation time of angular velocity. We assess natural variation in locomotor behavior and uncover persistent periods of preferred ground speed that correlate with the intensity of the odor response. We next develop a simple but biologically plausible computational model of locomotor control by a population of DN-like units, and show that graded changes in inhibition in this model can recapitulate the shifts in statistics that we observe experimentally. Finally, guided by connectome data, we gain genetic access to small populations of glutamatergic (putative inhibitory) neurons in the fly pre-motor center known as the Lateral Accessory Lobe (LAL) and test their role in shaping locomotor statistics. We find that activation of one population— labeling LAL089, LAL091, and LAL093— generates all three statistical signatures of offset search, while activation and silencing of this line bi-directionally modulates angular velocity at odor offset. A second group of neurons, including the single neuron pair LAL073, bi-directionally regulates walking speed, and can scale the length of the odor-evoked run. Our work highlights a simple computational architecture that can capture statistical features of locomotion across states and identifies potential neural correlates of this model.

## Results

### Locomotor statistics in walking flies vary both spontaneously and in response to sensory stimuli

In response to attractive odor, walking flies first run upwind, then perform a local search behavior that allows them to more readily re-locate a lost plume (Álvarez-Salvado *et al*, 2018, Demir et al. 2020). To understand how this local search behavior is generated, we analyzed ground speed and angular velocity in a previously collected set of behavioral trajectories (Álvarez-Salvado *et al*, 2018, see Methods). In these experiments, flies were placed in laminar wind tunnels and presented with a 10 second pulse of appetitive odor (10% apple cider vinegar, Fig. 1A-C). We examined the distribution of ground speeds and angular velocities in the 10 seconds following odor offset and compared these to baseline behavior prior to odor exposure (Fig. 1D-E, Supp Fig. 1A). Following odor offset, flies favor lower ground speeds and larger angular velocities (Fig. 1D-E), leading to higher angular dispersion (Supp Fig. 1B). During this offset period, flies also turned in the same direction for extended periods of time (Fig. 1C, arrows), leading to a wider autocorrelogram of the angular velocity (Fig. 1F). Turn frequency did not change significantly (Supp Fig. 1C). Three statistical features thus underlie the shift from baseline walking to local search: reduced ground speed, increased angular velocity, and longer history dependence in angular velocity.

**Fig 1:**
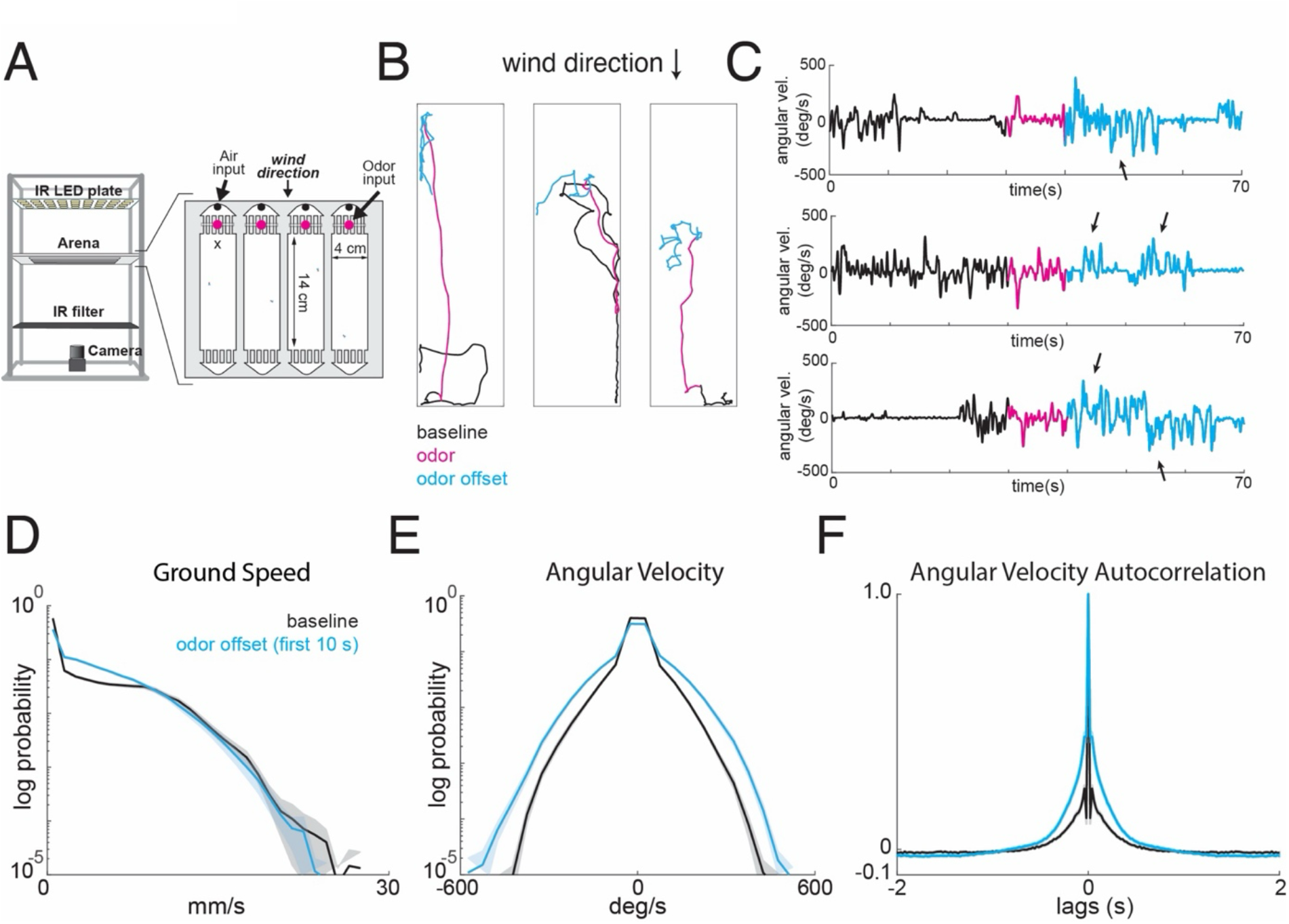
Odor loss drives systematic changes in locomotor statistics. A. Schematic of wind tunnel apparatus. Flies are constrained to walk in shallow 4 by 14 cm arenas and presented with temporally controlled wind and odor stimuli. Walking behavior is monitored using IR LEDs and a camera. B. Example trajectories of walking flies presented with a 10 s odor pulse (10% apple cider vinegar) centered in 70 s of laminar wind (left). Black: pre-odor baseline; magenta: odor period; cyan: post-odor. Flies move upwind towards the wind source in response to odor and perform a local search following odor offset. Wind direction indicated by arrow. C. Time course of angular velocity for examples shown in B. Note repeated turns in the same direction during post-odor period (cyan, arrows). D. Distribution of ground speeds during the post-odor search phase (first 10 s after odor, cyan) versus pre-odor baseline walking (first 10 s of baseline, black). Lower ground speeds are favored during the post-odor period compared to baseline. Error bars represent standard error across flies (n=1306 total trajectories from 75 flies). E. Distribution of angular velocities during the post-odor period (cyan) versus baseline walking (black). Larger angular velocities are favored during the post-odor period, compared baseline. Error bars represent standard error across flies. Same flies, trajectories, and time periods as in (D). F. Autocorrelation of angular velocity during the post-odor period (cyan) versus baseline walking (black). Autocorrelogram widens during the post-odor period. Error bars represent standard error across flies. Same flies, trajectories, and time periods as in (D).

We next examined spontaneous changes in locomotor behavior in trials without odor presentation. We observed periods of slow, local movement with many turns, and periods of quick, wide-spread movement with infrequent turns (Fig. 2A). We used a change point detection algorithm to segment spontaneous walking into epochs with different mean ground speeds (Fig 2B). The distribution of mean ground speeds had two peaks (Fig. 2C): a peak near zero (when flies were stopped), and a range of ground speeds from 5-20mm/s. Plotting mean angular velocity for each epoch as a function of mean ground speed revealed that angular velocity grows linearly with ground speed at low ground speeds (<5mm/s, arrow), but decreases at higher ground speeds (Fig. 2D). We additionally examined correlations between ground speed and angular velocity on the timescale of individual turns. We used a thresholding algorithm to extract turns, and sorted the associated behavioral segments by the mean ground speed in the half-second prior to the turn (Fig. 2E). At high ground speeds, we found that turns were associated with a brief drop in ground speed. This relationship is not present at low ground speeds. Very similar results were obtained during odor trials and odor periods (Supp Fig. 1D,E). Thus, ground speed and angular velocity exhibit complex interactions on multiple timescales.

**Fig 2:**
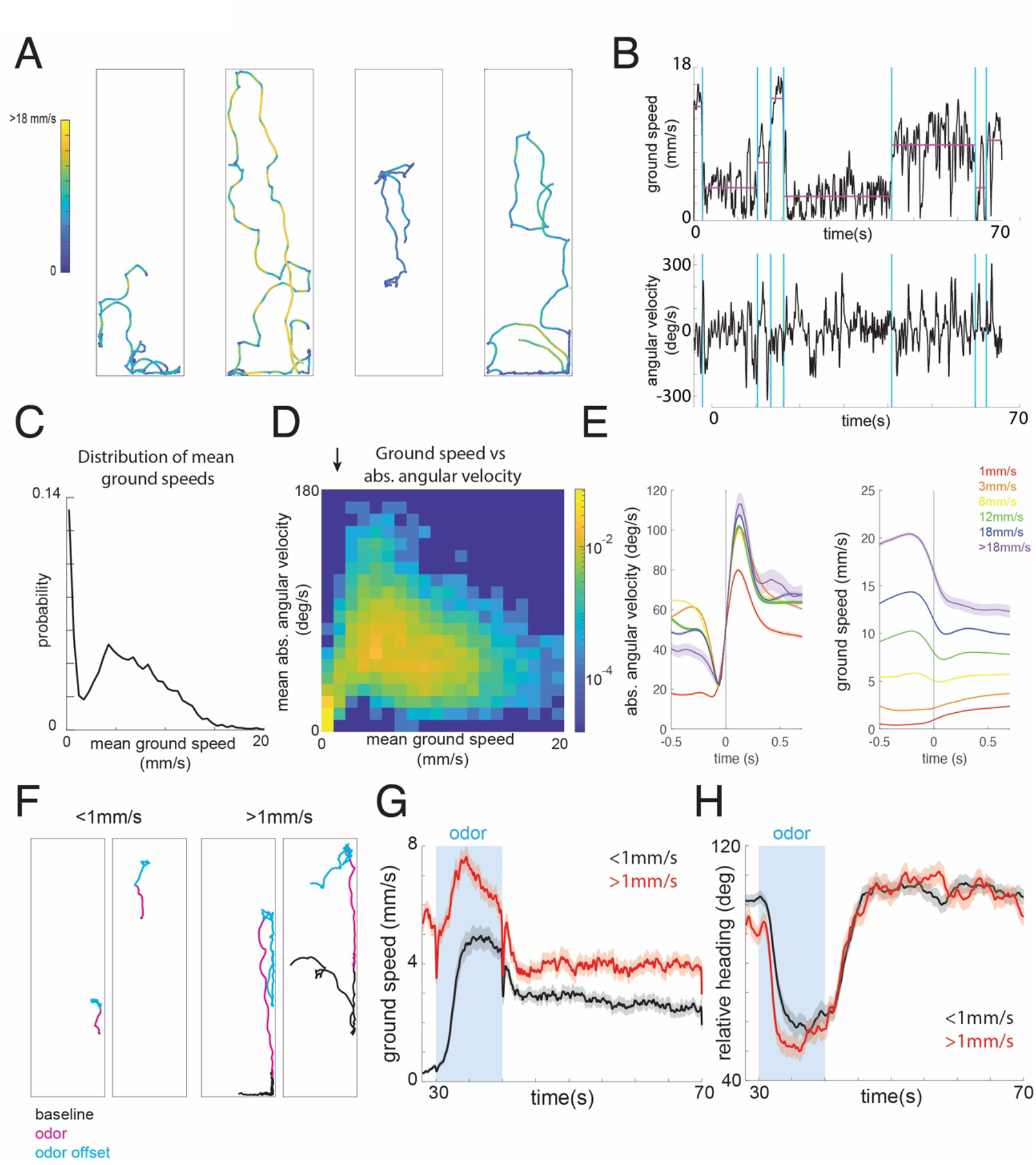
Spontaneous variation in locomotor states shapes responses to odor. A. Example trajectories illustrating spontaneous variation in locomotor statistics in the absence of odor. Colors indicate instantaneous ground speed as shown in color bar. B. Time courses of ground speed and angular velocity for rightmost trajectory in (A), segmented into behavioral epochs using change point detection. Blue vertical lines indicate epochs and purple lines represent mean ground speed for each epoch. C. Distribution of mean ground speeds during unstimulated walking for behavioral epochs segmented by change point detection, showing a bimodal distribution, corresponding to stationary and moving modes (n=898 trajectories from 56 flies). D. Joint distribution of mean ground speed and mean angular velocity for all epochs segmented by change point detection. Angular velocity grows with ground speed between ∼0-5 mm/s (black arrow), then decreases for higher ground speeds. Same flies as in (C). E. Angular velocity (left) and ground speed (right) relative to turn onset. Turns were detected by threshold crossings in angular velocity, then sorted by ground speed in the 500 ms prior to turn onset. Curves represent means within each ground speed range (n=898 trials from 56 flies, same flies as in C). Error bars represent standard error across turns within each category. At high ground speeds, we observe a decrease in ground speed preceding turn onset. At low ground speeds, acceleration follows turn onset. F. Example trajectories of flies moving <1 mm/s at odor onset (left) and flies moving >1 mm/s at odor onset (right). Flies that were moving >1 mm/s at odor onset travel further in response to the odor stimulus. G. Average ground speed for all trials where flies (N=75) were moving <1 mm/s at odor onset (black, 711 trials) and for trials where flies were moving >1 mm/s at odor onset (red, 595 trials). Trials where flies were moving >1 mm/s at odor onset show higher ground speeds during odor and after odor offset. Error bars represent standard error across fly means. H. Average relative heading (upwind direction at 0 degrees) for all trials where flies were moving <1 mm/s at odor onset (black) and for trials where flies were moving >1 mm/s at odor onset (red). Trials where flies were moving >1 mm/s at odor onset show a more rapid orientation to the upwind direction. Error bars represent standard error across fly means.

Finally, we asked whether the locomotor state of the fly can affect its response to a sensory stimulus. We sorted trials based on the mean ground speed prior to odor onset. Flies that were moving slowly prior to odor encounter produced shorter odor-evoked runs (Fig. 2F), with slower ground speeds during and after odor (Fig. 2G). The magnitude of the wind-orientation response was largely un-affected by prior ground speed, although the orienting response was slightly delayed (Fig. 2H). This analysis argues that low ground speed represents a behavioral state that can affect the response to odor input.

### A physiologically-inspired model of locomotor control

We next sought to develop a conceptual model of locomotor control that could account for the statistics we observed experimentally. Our model is intended to be relatively abstract, and to capture the statistical features of locomotion we describe above while retaining certain aspects of DN coding and physiology from the literature. Among these, we considered that (1) multiple DNs contribute to both forward and angular velocity (Rayshubskiy et al. 2024, Yang et al. 2023, Braun et al. 2024, Feng 2024), (2) different units make different contributions to forward versus angular velocity (Rayshubskiy et al. 2024, Yang et al. 2024, Bresovec et al. 2024, Aymanns et al, 2022), (3) bilateral activity correlates with forward velocity while activity differences between hemispheres correlate with angular velocity (Bidaye et al. 2020, Yang et al. 2023, Bresovec et al. 2024, Aymanns et al. 2022), and (4) distinct sets of DNs promote stopping (Lee and Doe 2021, Sapkal et al. 2024). Based on these considerations, we developed a simple model of locomotor control (Fig. 3A-C). In this model, each DN-like unit (*u*_1_-*u*_5_) can interact with the other units and makes a distinct contribution to overall locomotion (Fig. 3A,B). The *u*_3_ unit promotes stopping when active. In the absence of *u*_3_ activity, population activity across the remaining units drives locomotion. Forward velocity reflects a weighted sum of activity in the bilateral pairs *u*_1_, *u*_5_ and *u*_2_, *u*_4_, while angular velocity reflects a weighted sum of the differences in activity across these pairs. *u*_1_ and *u*_5_ have larger coefficients for forward velocity (α,β), while *u*_2_ and *u*_4_ have larger coefficients for angular velocity (γ,δ). Though highly simplified, this model captures the intuition that forward and angular velocity are regulated by graded activity across a population of DNs that can contribute differentially to these two parameters. The model coefficients (α,β,γ,δ) can be tuned to produce ground speed and angular velocity distributions similar to those observed experimentally (Supp. Fig 2A). By design, the model can reproduce the effects of activation in DNp09, in which unilateral activation produces ipsilateral turning, while bilateral activation produces an increase in ground speed (Bidaye et al. 2020, Supp Fig. 2B), and the effects of activating Pair1 or Brake neurons that drive stopping (Lee and Doe 2021, Sapkal et al. 2024, Supp Fig. 2C).

**Fig 3:**
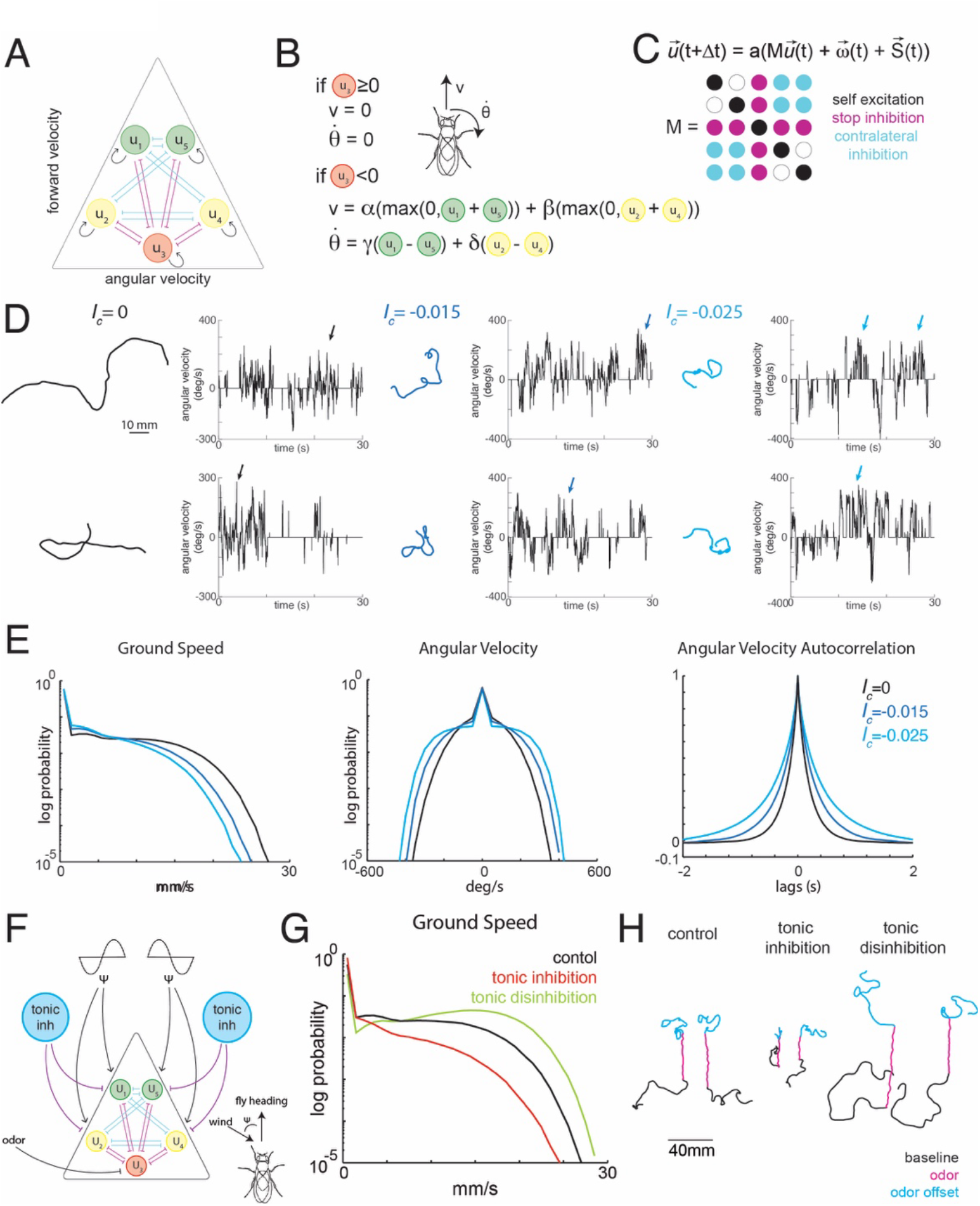
A physiologically inspired model can reproduce spontaneous and odor-evoked changes in locomotor statistics. A. Model locomotor control circuit comprised of five units. Green units (*u*_1_ and *u*_5_) drive large changes in forward velocity and small changes in angular velocity. Yellow units (*u*_2_ and *u*_4_) drive small changes in forward velocity and large changes in angular velocity. Red unit (*u*_3_) drives stopping. All units excite themselves (black arrows) through a uniform positive weight. Movement-producing units (*u*_1_, *u*_2_, *u*_4_, *u*_5_) and the stop unit (*u*_3_) mutually inhibit each other through a uniform inhibitory weight (stop inhibition, magenta). Left (*u*_1_ and *u*_2_) and right (*u*_4_and *u*_5_) movement-producing units mutually inhibit each other through a second uniform inhibitory weight (contralateral inhibition, cyan). B. Relationship between unit activity and locomotion. Positive activity in the stop unit (*u*_3_ ≥ 0) suppresses forward movement and turning. When activity in the stop unit is negative, forward velocity is calculated as a weighted rectified sum of all movement-producing units and angular velocity is calculated as a sum of the weighted difference between paired units on either side of the model. C. Unit dynamics are governed by a connectivity matrix *M*. Unit activity at time *t*+ Δ*t*is computed by multiplying the activity at the previous time step *t*by the matrix *M*, adding independent Gaussian noise 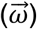 to each unit, as well as any inputs to the network 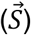 (see Methods), and then applying a sigmoid nonlinearity *a*. The network inputs 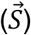 are the sum of the stop signal (eq. 5), the steering signal (eq. 6), and a feedforward tonic offset. See Methods for further details. D. Example trajectories of model flies with different amounts of contralateral inhibition (*I*_*c*_) and associated angular velocity traces. In the absence of contralateral inhibition (left), trajectories are relative straight and left and right turns are intermingled (black arrows). As contralateral inhibition increases, trajectories become more tortuous, and the model produces runs of turns in the same direction (blue and cyan arrows), similar to experimental trajectories observed during offset search. E. Statistical features of locomotion in model flies (from 1300 model trajectories) as a function of the strength of contralateral inhibition. Left: Ground speed distribution, center: angular velocity distribution, right: autocorrelation of angular velocity. Increased contralateral inhibition shifts the ground speed distribution towards lower values, shifts the angular velocity distribution towards higher values, and widens the autocorrelogram of angular velocity, similar to what we observe during post-odor search (Fig. 1D-F). F. Model schematic with the addition of odor- and wind-gated terms. To drive upwind running, odor suppresses the activity of the stop unit (*u*_3_) and modulates activity in the movement-producing units (*u*_1_, *u*_2_, *u*_4_, *u*_5_) according to a steering function that depends on wind direction. To modulate overall locomotor speed, tonic feedforward inhibition acts uniformly on all movement-producing units. G. Distribution of ground speeds produced by the model (from 500 model trajectories) with no feedforward inhibition (black), feedforward inhibition (red), and feedforward disinhibition (green). Tonic inhibition and disinhibition shift the distribution of ground speeds to lower or higher values respectively. H. Example odor-evoked trajectories produced by the model in the absence of feedforward inhibition (left), with feedforward inhibition (middle), and with feedforward disinhibition (right). Black: baseline walking; magenta: odor; cyan: post-odor search. Feedforward inhibition shrinks resultant trajectories, while disinhibition lengthens trajectories, mimicking shifts seen across behavioral modes in experimental data.

We next considered how activity arises dynamically in these DN-like units (Fig. 3C). Connectomic analysis suggests that different DNs promoting turning can receive input from largely non-overlapping upstream neurons (Yang et al. 2024). As a simplifying assumption, each unit in our model therefore receives an independent Gaussian noise input, ω_i_. Experiments in headless flies also suggest that DNs interact directly or indirectly within the brain to generate walking and turning behavior (Braun et al. 2024). Thus, our units can interact with themselves and each other through an interaction matrix M (Fig 3C). In our model, this matrix contains only three kinds of entries: a self-excitation term for each unit (black), bidirectional inhibitory interactions between the stop unit and all others (stop inhibition, magenta), and bidirectional inhibition between locomotor units in the two hemispheres (contralateral inhibition, blue). Activity in the units is limited by a sigmoidal activation function which prevents activity from growing without bound (see Methods).

Using this model, we asked if we could reproduce the shift between the statistics of baseline walking, and those of search behavior evoked by odor offset. With fixed parameters for self-excitation and stop inhibition, we found that increasing the magnitude of contralateral inhibition (*I*_*c*_) could shift the model from relatively straight trajectories resembling baseline walking to more tortuous trajectories resembling local search (Fig. 3D). In the absence of contralateral inhibition, left and right turns are intermingled (black arrows) resulting in generally straight and dispersive trajectories. In contrast, in the presence of contralateral inhibition, trajectories feature long runs of repeated turns in the same direction (blue and cyan arrows), similar to what we observe during search behavior evoked by odor offset. Quantitatively, increasing contralateral inhibition shifts the distribution of forward velocities to lower values, shifts the distribution of angular velocities to larger values, and widens the autocorrelation of angular velocity (Fig. 3E), similar to what we observed experimentally following odor offset. In the absence of contralateral inhibition, activity in locomotor units is correlated with each other, and anti-correlated with the stop unit, while in the presence of contralateral inhibition, units on the left and right become anti-correlated with one another (Supp. Fig. 2D), leading to larger angular velocities and lower forward velocities. Thus, a single parameter of the model governs the statistics of locomotor dispersal and smoothly switches behavior between baseline and search regimes.

Could the same effect be achieved by modulating ipsilateral connectivity? Several recent studies have highlighted ipsilateral excitation as an important motif underlying locomotor control (Braun et al. 2024, Feng et al. 2024). To explore this idea, we ran model simulations in which there were no contralateral interaction between locomotor units, but ipsilateral units were connected by either excitatory (Supp Fig. 3A-C) or inhibitory (Supp Fig. 3D-F) weights. We found that while ipsilateral excitation could shift the distribution of angular velocities to higher values and widen the angular velocity autocorrelogram, it also shifted the distribution of forward velocities to higher values (Supp Fig. S3C), resulting in trajectories that were dispersive, rather than localized (Supp Fig. S3B). Ipsilateral inhibition produced the opposite effect: a narrower distribution of angular velocities with a narrower autocorrelogram and lower groundspeeds. These effects arise because the ipsilateral models both lead to correlated activity in units on the two sides of the brain (Supp Fig. 3G,H), rather than anticorrelated activity, as seen with contralateral inhibition. Thus, in our simple model framework, only contralateral inhibition is capable of reproducing the reduction in ground speed and increase in angular velocity seen during offset search.

We next asked if the model could reproduce the effects of baseline groundspeed on odor responses. To generate an odor response, we incorporated two odor-gated terms into our model (Fig 3F): a steering function that depends on the angle of the fly with respect to the upwind direction, and an odor-driven suppression of the stop unit (*u*_3_). The steering function might represent the output of PFL3 steering neurons in the central complex (Matheson et al. 2022, Westeinde et al. 2024, Mussels Pires et al. 2024), which are known to impinge on descending neurons controlling steering (Westeinde et al. 2024). These additions produce an upwind run during odor, while increasing contralateral inhibition can then produce search behavior at odor offset (Supp. Fig 2E). We then added feed-forward inhibition that controls the activity of all locomotor units (Fig. 3F). Strikingly, increasing or decreasing feed-forward inhibition caused both a shift in distribution of ground speeds (Fig. 3G), and caused the odor-evoked run to shrink or grow, similar to what we observed experimentally during spontaneous behavioral states (Fig. 3H). Thus, the conceptual architecture we describe here, in which DN-like units receive both steering and speed control signals and jointly regulate forward and angular velocity, allows a state-dependent signal to scale the response to a sensory stimulus, as observed in our experimental data.

An advantage of this simple conceptual framework is that it is amenable to mathematical analysis, which allows us to understand its behavior. As described in the Mathematical Appendix, the dynamics of the units can be decomposed into two subsystems through a linear transformation of coordinates: a symmetric subsystem that controls forward velocity, and an anti-symmetric subsystem that controls angular velocity. The inputs to the units, such as the steering commands and feedforward inhibition, can be expressed in this new coordinate system, where it is clear that steering only impacts the anti-symmetric subsystem, while feed-forward inhibition only impacts the symmetric subsystem. This explains how two multiplexed signals can independently control orientation and speed.

This analysis also allows us to understand why increasing contralateral inhibition shifts the model from relatively straight trajectories to more tortuous ones. The dynamics of the anti-symmetric subsystem (which controls turning) depend on its eigenvalues; when these are all negative, the model exhibits stable fixed-point dynamics, and activity returns to zero— the only stable point under these conditions — following an input. In this regime, turns to the left and right are interspersed, and the model produces mostly straight trajectories (Supp. Fig. 4, white background). In contrast, when the anti-symmetric system has a positive eigenvalue, the fixed point at zero becomes unstable and tends to go towards one of the saturation points, until noise pushes it back towards zero. Under these conditions, the model produces runs of repeated turns in the same direction, resulting in more tortuous looping trajectories (Supp. Fig. 4, gray background). The duration of these runs increases with the strength of contralateral inhibition, and depends on the amount of instability, as well as on the characteristics of the noise.

**Fig 4:**
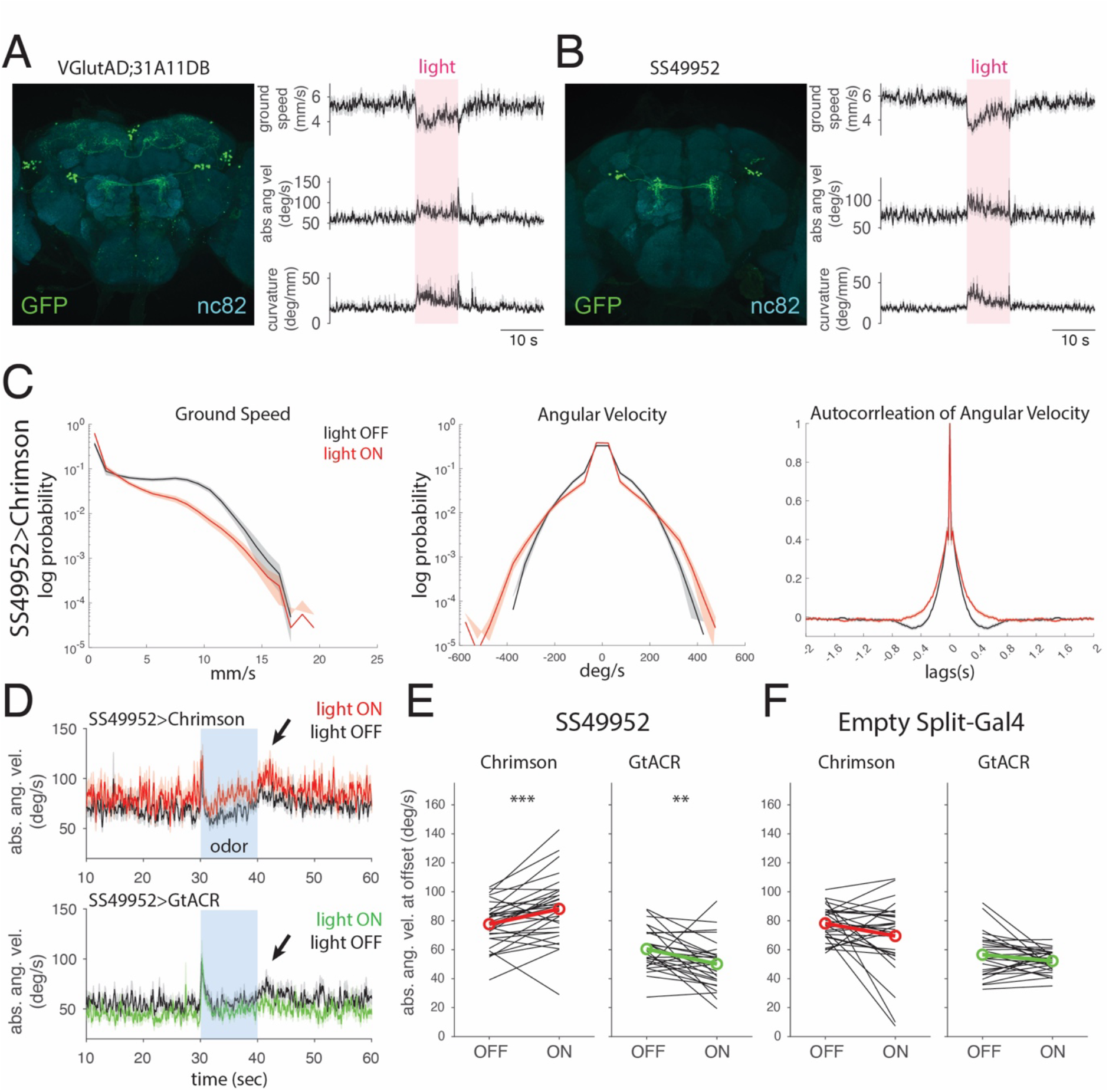
Glutamateric neurons that regulate curvature. A. (Left) Confocal image of neurons labeled by VGlut-AD;31A11-DBD > GFP (green). Neuropil (nc82) in cyan. This line labels 10-11 neurons in each LAL that project contralaterally to the opposite LAL, as well as populations of neurons in SIP and LH and in the VNC (see Supp. Fig 4A). (Right) Average time course of forward velocity, angular velocity, and curvature response to Chrimson activation in VGlut-AD;31A11-DBD neurons (red: 10s light). Error bars represent standard error across fly means (n=183 total trials from 31 flies). Activation causes a decrease in ground speed, an increase in angular velocity, and an increase in curvature. B. (Left) Confocal image of neurons labeled by SS49952 > GFP (green). Neuropil (nc82) in cyan. This line labels 8-9 neurons in each LAL that project contralaterally to the opposite LAL, as well as 1-2 neurons in the VNC (see Supp. Fig 4A). (Right) Average time course of forward velocity, angular velocity, and curvature response to Chrimson activation in SS49952 neurons (red: 10s light). Error bars represent standard error across fly means (250 total trials from 31 flies). Activation causes a decrease in forward velocity, an increase in angular velocity, and an increase in curvature. C. Distributions of ground speed (left), angular velocity (middle), and angular velocity autocorrelation (right) in SS49952 > Chrimson flies in the absence (black) and presence (red) of light. Error bars represent standard error across flies (n=250 total trials from 31 flies). Activation shifts ground speed towards lower values, shifts angular velocity towards higher values, and widens the angular velocity autocorrelation, similar to post-odor search (Fig. 1D-F). D. (Top) Average time course of absolute angular velocity in SS49952 > Chrimson flies responding to odor (blue: 10s) in the absence (black) and presence (red) of light. Activation causes an increase in angular velocity during the post odor offset search, indicated by the black arrow (n=305 trials from 31 flies). (Bottom) Average time course of angular velocity in SS49952 > GtACR flies responding to odor (blue: 10s) in the absence (black) and presence (green) of light. Silencing of SS49952 neurons causes a decrease in angular velocity during the post odor offset search (n=313 total trials from 27 flies). E. Mean absolute angular velocity of SS49952 > Chrimson and SS49952 > GtACR flies in the absence (OFF) and presence (ON) of light during the post odor search phase (first 5 s after odor stimulus ends). Activation causes a significant increase in angular velocity during the post odor phase (paired *t*-test, p=9.35e-04, n=350 total trials from 31 flies). Silencing causes a significant decrease in angular velocity during the post odor phase (paired *t*-test, p=0.0093, n=313 total trials from 27 flies). Black lines: average data from individual flies. Red and green lines: group averages. F. Mean absolute angular velocity of empty split-Gal4 > Chrimson and empty split-Gal4 > GtACR flies in the absence and presence of light during the post odor search phase (first 5 s after odor stimulus ends). Activation causes no significant changes in angular velocity during the post odor phase (paired *t*-test, p=0.0672, n= 281 total trials from 32 flies). Silencing causes no significant changes in angular velocity during the post odor phase (paired *t*-test, p=0.0900, n=302 total trials from 30 flies). Black lines: average data from individual flies. Red and green lines: group averages.

The eigenvalues of the anti-symmetric system depend on both the self-excitation parameter and on contralateral inhibition (Supp. Fig. 4). However, we find the that the model is most flexible when self-excitation is set at the transition point from stable to unstable behavior (Supp. Fig. 4, middle row). In this regime, small changes in contralateral inhibition smoothly shift the model from relatively straight to relatively tortuous trajectories. When self-excitation places the model mostly in the stable regime (top row), much larger values of contralateral inhibition are required to generate search-like trajectories, while when self-excitation places the model in the unstable regime (bottom row), trajectories are unrealistically tortuous even without contralateral inhibition. For this reason, for the simulations shown in Fig. 3, we have set the self-excitation term to this transition point. Our model and analysis suggest that a locomotor network poised between stable and unstable regimes may allow an animal to smoothly regulate the tortuosity of its trajectories through graded changes in inhibition between the two hemispheres. The Appendix analysis also shows how inhibitory contralateral connectivity allows for modulation of forward velocity and angular velocity in opposite directions (Figure 3E), while a model with only ipsilateral connectivity co-regulates forward and angular velocity (Figure S3C,F).

Finally, we show analytically that the model exhibits similar dynamics regardless of the number of locomotion-promoting units on each side, so long as the structure of the interaction matrix M is maintained. In particular, the role of contralateral inhibition in governing the tortuosity of trajectories remains the same, regardless of the number of units. In the fly brain, multiple DN types have been identified that show activity correlated with turning (Yang et al. 2023, Aymanns et al. 2022) and the number of such units likely varies across species (Okada et al. 2003, Staudacher 1998). Thus, the analysis presented here, and the proposed role of contralateral inhibition in governing tortuosity, could apply to real biological networks with variable numbers of locomotor control units.

### Glutamatergic premotor neurons provide complementary control of locomotor statistics

Our model and analysis suggest that inhibition— in particular contralateral inhibition between the two hemispheres— plays a key role in shaping locomotor statistics. Connectomics (Scheffer et al. 2020, Greg Jefferis, personal communication) reveals that the lateral accessory lobe (LAL), a pre-motor center strongly implicated in steering (Kanzaki et al. 1994, Namiki and Kanzaki, 2016), contains several populations of decussating neurons that are predicted to express glutamate (Eckstein et al. 2023), often an inhibitory transmitter in the central fly brain (Liu and Wilson 2013). We therefore sought to gain genetic access to these populations.

By combining a VGlut AD hemidriver with the hemidriver 31A11 DB, we generated a line labeling contralaterally projecting neurons in the LAL (Fig. 4A), as well as other populations in the brain and ventral nerve cord (VNC, Fig. 4A, Supp Fig. 5A). Activating this line with Chrimson drove a decrease in ground speed and an increase in angular velocity, resulting in an increase in curvature (Fig. 4A, Supp Fig. 5C), similar to behavior during offset search. We therefore named this line Tortuous-Gal4. Seeking a cleaner line labeling the same LAL neurons, we identified SS49952 (24H08AD;31A11DB) which labels 8-9 LAL neurons and 1-2 neurons in the VNC (Fig. 4B, Supp. Fig. 5A,B. Activating SS49952 drove a similar but stronger phenotype to Tortuous-Gal4 (Fig. 4B,C, Supp Fig 5C), including all three signature of offset search: a shift of the groundspeed distribution towards lower values, a shift of the angular velocity distribution towards higher values, and a widening of angular velocity autocorrelation (Fig. 4C). To test the role of these neurons in odor-evoked offset search, we compared responses of flies to an odor pulse (1% ACV) in interleaved trials with and without SS49952 activation or silencing (Fig. 4D). Activating SS49952 neurons increased angular velocity during offset search, while silencing SS49952 neurons decreased angular velocity during offset search (Fig 4D,E). These effects were not observed when Chrimson or GtACR were driven by empty-Gal4 or empty split-GAL4 drivers (Fig 4F, Supp Fig. 5C,D). Silencing of Tortuous-Gal4 or SS49952 with GtACR had no significant effect on motor output in the absence of odor (Supp Fig. 5D), suggesting that these neurons do not contribute strongly to baseline locomotion.

**Fig 5:**
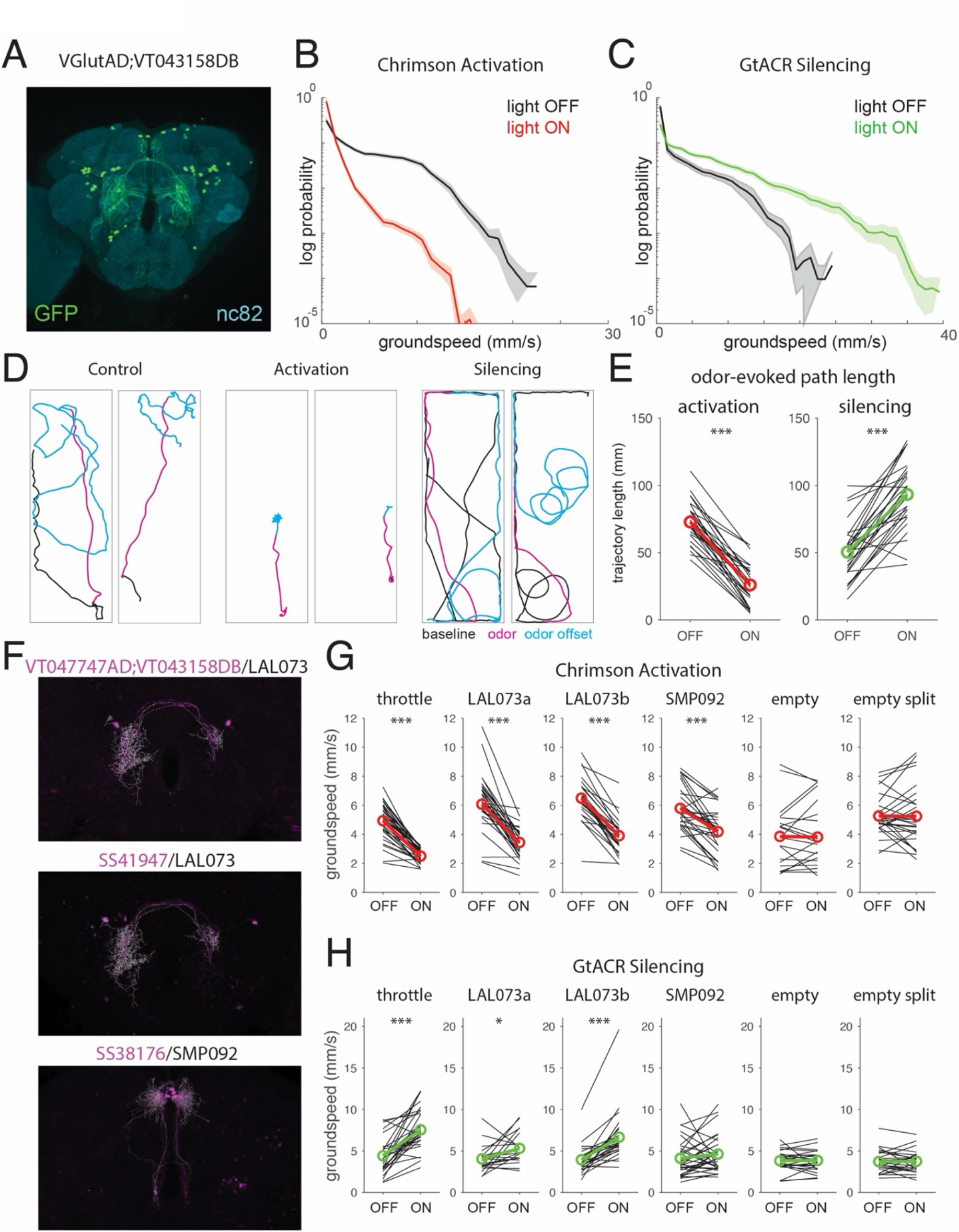
Glutamatergic neurons that regulate ground speed. A. Confocal image of neurons labeled by VGlut-AD;VT043158-DBD > GFP (green). Neuropil (nc82) in cyan. This line labels 14-15 neurons in each LAL that project contralaterally to the opposite LAL, 2 bilateral pairs of neurons along the dorsal midline, as well as other neurons in the brain and VNC (Supp. Fig. 5A). B. Distribution of groundspeeds of VGlut-AD;VT043158-DBD > Chrimson flies in the presence of light (red) and absence of light (black). Traces represent mean ± standard error across flies (n=337 total trials from 34 flies). Activation shifts groundspeeds to lower values. C. Distribution of groundspeeds of VGlut-AD;VT043158-DBD > GtACR1 flies in the presence of light (green) and absence of light (black). Traces represent mean ± standard error across flies (n=219 total trials from 23 flies). Silencing shifts groundspeeds to higher values. D. Example trajectories in response to an odor pulse (10s of 1% ACV) in the absence of light (Chrimson control, left), during optogenetic activation of VGlut-AD;VT043158-DBD neurons with Chrimson (middle), and during optogenetic silencing of VGlut-AD;VT043158-DBD neurons with GtACR1 (right). E. Path length of odor-evoked runs under different stimulation conditions. Activation of VGlut-AD;VT043158-DBD neurons significantly reduces path length (paired *t*-test, p= 8.4681e-15, 192 total trials from 28 flies) while silencing significantly increases path length (paired *t*-test, p= 1.1892e-09, n=229 total trials from 29 flies). Black lines: average data from individual flies. Red and green lines: group averages. Asterisks: statistically significant changes in paired *t*-tests. F. Warped confocal images of neurons labeled by VT047747AD;VT043158DB, SS41947, and SS38146 > GFP (magenta) overlaid with best-fit neurons from the hemibrain connectome (white). VT047747AD;VT043158DB labels LAL073, SS41947 labels LAL073, and SS38146 labels SMP092. G. Mean groundspeeds of each line expressing Chrimson in the absence (OFF) versus presence (ON) of light. Black lines: average for each fly. Red lines: average across flies. Asterisks: statistically significant changes in paired *t*-tests. Throttle: p = 7.6288E-15, 337 total trials from 34 flies. VT047742;VT043158DB: p = 1.5877E-10, 198 total trials from 34 flies. SS41947: p = 4.4820E-12, 263 total trials from 28 flies. SS38176: p = 9.5804E-07, 215 total trials from 30 flies. Empty-Gal4: p = 0.51294, 411 trials from 25 flies. Empty split-Gal4: p = 0.94536, 146 trials from 30 flies. H. Mean groundspeeds of each line expressing GtACR in the absence (OFF) versus presence (ON) of light. Black lines: average for each fly. Red lines: average across flies. Asterisks: statistically significant changes in paired *t*-tests. Throttle: p = 2.5372E-07, 219 total trials from 31 flies. VT047747AD;VT043158DB: p = 1.5625E-02, 121 total trials from 26 flies. SS41947: p = 7.6295E-06, 230 total trials from 29 flies. SS38176: p = 0.28702, 179 trials from 31 flies. Empty-Gal4: p = 0.69946, 224 trials from 29 flies. Empty split-Gal4: p = 0.97136, 227 trials from 30 flies.

Anatomical analysis suggests that the most likely LAL neurons labeled by SS49952 are LAL089,LAL091, and LAL093 (Supp Fig. 5B) which are all predicted to be glutamatergic (Eckstein et al. 2024), consistent with our genetic labeling strategy. These neurons receive a small input from PFL2 output neurons of the Central Complex (Supp Fig. 5E), which drives slowing and a non-directional increase in turning when activated (Westeinde et al. 2024). These data suggest that putative inhibitory neurons within the brain can modulate locomotor statistics and odor offset behavior in the manner predicted by our model, although it remains possible that some of the effects of SS49952 manipulation could arise from the 1-2 VNC neurons labeled in this line.

Combining the VGlut AD hemidriver with a second hemidriver, VT043158 DB, we generated a line labeling a different population of contralaterally-projecting LAL neurons (Fig. 5A), small clusters of neurons in the superior medial protocerebrum (SMP), and several clusters in the VNC (Supp Fig. 6A). Activating this line with Chrimson strongly decreased ground speeds (Fig. 5B,G), while silencing this line with GtACR increased ground speeds to an atypical range (Fig. 5C,H). We therefore named this line Throttle-Gal4. Combining optogenetic manipulation of this line with odor stimulation, we found that activation of Throttle-Gal4 decreased the length of upwind runs while silencing lengthened these runs (Fig. 5D,E), similar to the predictions of our model for feedforward inhibition.

To identify the neurons responsible for these phenotypes, we made or obtained three cleaner split-Gal4 lines containing the same VT043158 hemidriver (Fig. 5F). Two of these lines— VT047747AD;VT043158DB, and SS41947— both label the single neuron pair LAL073 with extremely sparse and non-overlapping VNC expression (Supp Fig. 6A). Activating each line produced a decrease in mean groundspeed (Fig. 5G) and a shift in the groundspeed distribution to lower values (Supp. Fig. 6B), while silencing increased mean groundspeed (Fig. 5H) and shifted the groundspeed distribution to higher values (Supp Fig. 7A). These effects were similar to, but weaker than, the effects of manipulating throttle-Gal4 (Fig. 5G,H, Supp Fig. 6B,C, 7A,B). A third line, SS38176, labels 4 SMP092 neurons in the dorsal SMP, with very sparse VNC expression (Supp Fig. 6A). Activating this line reduced groundspeed (Fig. 5G, Supp Fig. 6B,C) but silencing had no significant effect on behavior (Fig. 5H, Supp Fig. 7A,B). Each of these neurons is predicted to be glutamatergic (Eckstein et al. 2024) and none had any effect on angular velocity (Supp Fig. 6D,7C).

Together these data suggest that Throttle-Gal4 labels multiple neurons that regulate forward velocity, with LAL073 playing a prominent, but not exclusive, role in this effect, and SMP092 and other neurons contributing additively or synergistically. LAL073 receives its dominant input from PFL2, with additional input from several non-canonical MBONs (Supp Fig. 7D), and a very small input from PFL3 neurons, which form the steering outputs of the Central Complex (Stone et al. 2017, Westeinde et al. 2024, Mussells Pires et al. 2024). Thus, LAL073 may contribute to the slowing evoked by PFL2 activation. Together, these data provide experimental support for the hypothesis that distinct types of pre-motor inhibition can regulate groundspeed and tortuosity in a complementary and independent manner.

## Discussion

### Statistical analysis of odor-evoked locomotor behavior

During foraging, many animals respond to food odor cues by altering the statistics of their locomotion to search near one location, or to disperse more widely (Shtonda and Avery 2006, Ben Arous et al 2009, Davies et al 2015, Steck et al 2013, López-Cruz et al 2019, Leitch et al. 2021). Fruit flies have been shown to respond to the loss of an attractive odor with head casting in larvae (Gomez-Marin et al. 2011), local search when walking (Alvarez-Salvado et al. 2018, Demir et al. 2020), and casting or circling behavior in flight (van Breugel et al. 2014, Stupski et al. 2024). Previous studies have described walking local search as an increase in turn frequency (Alvarez-Salvado et al. 2018, Demir et al. 2020). Here we show instead that it arises from systematic shifts in the distribution and correlation of locomotor parameters.

We further explored variation in odor-evoked behavior and found that the length of the odor-evoked run depends on baseline groundspeed. We observed an effect of baseline groundspeed in spontaneous behavior and could accentuate this by experimentally manipulating groundspeed using our newly-discovered Throttle-Gal4 line. In contrast, upwind turning behavior was slightly slower in non-moving flies, but reached the same endpoint. These results argue that speed and direction can be encoded separately to produce an upwind run during odor.

### A modeling framework for population control of locomotion

Previous models of walking behavior in *Drosophila* have used variations of Hidden Markov Models (HMMs, Katsov et al 2017, Calhoun et al 2019, Tao et al 2019). These models rely on behavioral segmentation and classification and assume that walking can be decomposed into a series of discrete motifs. However, walking behavior in flies evolves on longer timescales that cannot be captured through a pure Markovian process (Berman et al 2016). Nested HMMs and those with dynamic transition matrices take steps towards capturing the multiple timescales within walking behavior (Calhoun et al 2019, Tao et al 2019). Non-Markovian models of fruit fly walking, on the other hand, more accurately capture long-term shifts in behavior caused by odor stimuli (Alvarez-Salvado et al 2018, Demir et al 2020) but fail to accurately capture natural distributions of forward and angular velocity. While current models of fruit fly walking capture different aspects of behavioral dynamics, few provide insight into the underlying neural structures that govern locomotor statistics.

Here we developed a simple model framework for continuous control of locomotion by a population of DN-like units. By design, our model is highly simplified, while retaining some aspects of biological encoding. The key aspects of this model are: (1) each unit makes a unique contribution to forward and angular velocity such that these parameters are controlled by activity across the populations a whole, and (2) units interact through a connectivity matrix that allows activity in one unit to influence the others. This simple model generates realistic distributions of forward velocity, angular velocity, and angular velocity correlations in a variety of behavioral states, and implicates inhibition in regulating these distributions to control tortuosity and speed. The importance of inhibition to our model led us to identify new neural populations that modulate tortuosity and speed.

In our simple model framework, where the same units control forward and angular velocity through their sums and differences, we found that ipsilateral connectivity could not produce the observed differences between baseline walking and offset search. However, recent studies have identified single units that primarily encode turning with little contribution to forward velocity (Yang et al. 2024, Feng et al. 2024). If turning and forward velocity are controlled separately, then ipsilateral excitation should be able to promote more prolonged sequences of turns, as has recently been suggested (Feng et al. 2024).

The simplicity of our model allows for rigorous mathematical analysis allowing insight into its function. For example, we find that our model is most flexible when its parameters position its dynamics at the edge of instability. In the stable regime, fluctuations in internal signals rapidly return to zero, so there are few sustained periods of turning in one direction or the other, even in the absence of a stabilizing external “goal” signal. In the unstable regime, fluctuations can push activity towards extreme values, where they remain until rare, large noise fluctuations cause them to return to zero, resulting in prolonged turning in one direction. When the model is positioned at the border between these two regimes, graded increases in inhibition can smoothly increase the propensity to instability and prolonged turning. This prediction could be tested by coupling population imaging from DNs with optogenetic perturbations.

In addition, we find analytically that the dynamics of the model do not depend on the number of units per side; rather, the critical feature is that units activity can be decomposed into symmetric (speed controlling) and asymmetric (turn controlling) subsystems. This suggests that similar dynamics and control principles might be found in a variety of nervous systems with different numbers of descending locomotor control neurons or with more segregated control of speed and turning. In the future, we envision that our model framework— a matrix of interactions between locomotor control units, with a separate set of downstream equations that describe the transformation from unit activity into locomotor features— could be expanded to include realistic numbers of DNs (Namiki et al. 2018), specific connectivity patterns drawn explicitly from *Drosophila* connectomes (Scheffer et al. 2020, Cheong et al. 2024, Braun et al. 2024, Sapkal et al. 2024), more realistic control of turning maneuvers (Yang et al. 2024, Feng et al. 2024), or correlated noise input to locomotor units (Braun et al. 2024). Such models would allow for a principled understanding of how connectivity in a population of locomotor control units affects the dynamics of locomotion across timescales.

### Identification of neural populations controlling walking tortuosity and speed

The lateral accessory lobe (LAL) has long been associated with turning behavior in insects (Kanzaki et al. 1994, Namiki and Kanzaki, 2016) and houses descending and local neurons whose activity correlates with and drives turning (Rayshubskiy et al. 2024, Yang et al. 2024, Feng et al. 2024). The LAL is also a major target of outputs of the Central Complex that drive goal-directed steering (Stone et al. 2017, Hulse et al. 2020, Mussels Pires et al. 2024, Westeinde et al. 2024). Inspired by our model, we investigated whether contralaterally projecting neurons in the Lateral Accessory Lobe (LAL) could generate the statistical changes in locomotion predicted by our model. We identified the line SS49952 (LAL089, LAL091, and LAL093) that produces all three statistical signatures of offset search when activated, and can bi-directionally regulate angular velocity at odor offset. We identified a second neuron— LAL073— that strongly and bi-directionally regulates walking speed. We also identified a third population— SMP092— whose activation decreases groundspeed. The dorsal SMP has been implicated in speed control by pan-neuronal imaging (Bresovec et al. 2024) and genetic analysis (Bidaye et al. 2020). As the effects of manipulating LAL073 or SMP092 were weaker than the full throttle-Gal4 line, it is likely that walking speed is encoded by a population of neurons in both the LAL and SMP. In flying flies, a large population of closely related neurons known as DNg02 regulate flight speed through a population code (Namiki et al. 2022). Our results are consistent with a population code for control of walking speed as well.

While our genetic analysis revealed populations that are potential neural correlates of our model, several caveats remain. First, while the line SS49952 produced similar phenotypes to offset search, it also labeled 1-2 VNC neurons. As we were not able to find a second line labeling these same LAL neurons with different VNC expression, it remains possible that the behavioral phenotypes arise from VNC neurons labeled in this line. Second, while our genetic strategy using VGlut AD, as well as connectomic predictions of neurotransmitter identity, suggest that our neurons were glutamatergic, we have not confirmed this functionally. Glutamate is often an inhibitory transmitter in the central brain (Liu and Wilson, 2013), but recordings from downstream partners will be needed to directly test whether these neurons are functionally inhibitory. Finally, imaging or recording from behaving flies will be required to conclusively link the neural populations identified here to search behavior and walking speed.

In vertebrates, distinct areas of the midbrain and brainstem have been shown to regulate speed and turning behavior across species. The mesencephalic locomotor region (MLR) can initiate and control locomotor speed across terrestrial, aquatic, and avian species (Shik et al.

1966, Kashin et al. 1974, Sirota et al. 2008, Steeves et al. 1987). Similarly, in both mice and zebrafish, a homologous set of reticulospinal neurons labelled by chx10 drive turning through ipsilateral modulation of oscillatory spinal circuits (Cregg et al 2018, Orger et al 2008, Huang et al 2013, Kimura et al 2006). How speed and turning are jointly regulated to produce area-restricted search and dispersal behaviors has been most studied in invertebrate species. In larval Drosophila, SEZ neurons have been implicated in modulating the rate of transition between runs and turns (Tastekin et al. 2015). In *c. elegans*, local search is initiated when glutamatergic signaling suppresses activity in two neurons that inhibit local search, AIA and ADE (López-Cruz et al. 2019). In flying flies, a large inhibitory interneuron known as VES041 has recently been identified that suppresses saccades, suggesting it may play a role in shifting behavior from local search to dispersal (Ros et al. 2023). Our analysis, model, and identification of neurons controlling curvature and walking speed provide a basis for asking whether neural circuits for area-restricted search exhibit conservation across diverse body plans and modes of locomotion.

## Acknowledgements

The authors would like to thank Greg Jefferies and Salil Bidaye for helpful suggestions about neural correlates of our models, Alex Bates for assistance with warping of confocal images and alignment to hemibrain neurons, and Matthew Clark, Claude Desplan, Tom Clandinin, and the Bloomington Drosophila Resource Center for fly lines. Emily Hao assisted with a subset of behavioral experiments. Matthieu Louis, Shy Shoham, and Christina Savin, as well as members of the Nagel lab provided feedback on the manuscript. This work was funded by F31 DC019553 to H.G. and R01DC107979 and RF1NS127129 to K.I.N., as well as an NSF neuronex grant (Odor2Action: NSF 2014217) to K.I.N., G.B.E., and J.D.V.

## Data and Code Availability

All data and code generated during this study will be made publicly available on Github and Zenodo upon publication.

## Materials and Methods

### Fly Stocks and Culture

All experimental flies were raised at 25°C on standard cornmeal-agar medium, with a 12h light-dark cycle. All experiments were performed on female flies between 3-8 days old. Genotypes used in each figure were as follows:

|  |  |
| --- | --- |
| Figure 1-2,<br>Fig. S1 | *W<sup>1118</sup> norpA<sup>36</sup>* |
| Figure 4 | A: VGlut-AD; 31A11-DBD > UAS-GFP, VGlut-AD; 31A11-DBD > UAS-Chrimson(III)<br>B,C: SS49952 > UAS-GFP, SS49952 > UAS-Chrimson(III)<br>D,E: SS49952 > UAS-Chrimson(III), SS49952 > UAS-GtACR(III)<br>F: empty Split-Gal4 > UAS-Chrimson(III), empty Split-Gal4 > UAS-GtACR(III) |
| Figure S4 | A: VGlut-AD; 31A11-DBD > UAS-GFP, SS49952 > UAS-GFP<br>B: SS49952 > UAS-GFP<br>C: VGlut-AD; 31A11-DBD > UAS-Chrimson(III)<br>SS49952 > UAS-Chrimson(III)<br>empty-Gal4 > UAS-Chrimson (III)<br>empty Split-Gal4 > UAS-Chrimson(III)<br>D: VGlut-AD; 31A11-DBD > UAS-GtACR(III)<br>SS49952 > UAS-GtACR(III)<br>empty-Gal4 > UAS-GtACR(III)<br>empty Split-Gal4 > UAS-GtACR(III) |
| Figure 5 | A: VGlut-AD; VT043158-DBD > UAS-GFP,<br>B,D,E: VGlut-AD; VT043158-DBD > UAS-Chrimson(III)<br>C,D,E: VGlut-AD; VT043158-DBD> UAS-GtACR1(III)<br>F: VT047747AD;VT043158DB>UAS-GFP<br>SS41947 > UAS-GFP<br>SS38176 > UAS-GFP<br>G: VGlut-AD; VT043158-DBD > UAS-Chrimson(III)<br>VT047742AD;VT043158DB > UAS-Chrimson(III)<br>SS41947 > UAS-Chrimson(III)<br>SS38176 > UAS-Chrimson(III)<br>empty-Gal4 > UAS-Chrimson (III)<br>empty Split-Gal4 > UAS-Chrimson(III)<br>H: VGlut-AD; VT043158-DBD > UAS- GtACR (III)<br>VT047742AD;VT043158DB > UAS- GtACR (III)<br>SS41947 > UAS- GtACR (III)<br>SS38176 > UAS- GtACR (II)<br>empty-Gal4 > UAS- GtACR (III)<br>empty Split-Gal4 > UAS- GtACR (III) |
| Figure S5 | A: VGlut-AD; VT043158-DBD > UAS-GFP<br>VT047747AD;VT043158DB>UAS-GFP<br>SS41947 > UAS-GFP<br>SS38176 > UAS-GFP<br>B,C,D: VGlut-AD; VT043158-DBD > UAS-Chrimson(III)<br>VT047747AD;VT043158DB > UAS-Chrimson(III)<br>SS41947 > UAS-Chrimson(III)<br>SS38176 > UAS-Chrimson(III)<br>empty-Gal4 > UAS-Chrimson (III)<br>empty Split-Gal4 > UAS-Chrimson(III) |
| Figure S6 | A,B,C: VGlut-AD; VT043158-DBD > UAS- GtACR (III)<br>VT047747AD;VT043158DB > UAS- GtACR (III) |
|  | SS41947 > UAS- GtACR (III)<br>SS38176 > UAS- GtACR (II)<br>empty-Gal4 > UAS- GtACR (III)<br>empty Split-Gal4 > UAS- GtACR (III) |

Experimental flies were collected at least 1 day post eclosion and kept at room temperature in custom made, time-shifted, light boxes for 2-7 days to acclimate to temperature and circadian rhythm. For optogenetic activation, 50 μl of 35mM all-trans retinal stock was mixed into ∼1 tsp of hydrated potato flakes and added to standard food vials at least four days prior to the experiment. All behavioral experiments were performed between subjective ZT0-ZT4. Flies were starved for ∼24 hours before the start of the experiment in empty plastic vials with a dampened Kimwipe for hydration.

A subset of analysis presented here (Figures 1 and 2) use a previously collected dataset of behavioral data from wind tunnel experiments completed using 10% apple cider vinegar. Collection and earlier analysis of these data are described in Alvarez-Salvado et al. 2018 (see Key Resources Table) but followed the same protocols described here.

### Wind tunnel experiments

Behavioral experiments in freely-walking flies were performed in miniature wind tunnel arenas as described previously (Álvarez-Salvado *et al*, 2018). Flies were placed in shallow 4cm by 14cm arenas that constrained them to walk. Wind was constant at ∼12cm/s. The arenas were backlit with IR LEDS (850nm, Environmental Lights) and monitored from below with a camera (Basler acA1920-155um). Stimuli were controlled through a NIDAQ board. Position and orientation data were collected in real time using custom LABVIEW code.

Flies were run for approximately 2 hours during which they were exposed to 6 randomly interleaved trial types: a blank control trial, a 10 second pulse of 1% apple cider vinegar, light on for the entire 70 seconds, a 10 second light pulse centered in the 70 second trial, a simultaneous 10 s pulse of odor with a 10s pulse of light, and a 10s odor pulse with light on for the entire 70 seconds. Each trial lasted 70 seconds with ∼5 seconds between trials. For Chrimson experiments, red light was presented using red LED strips (NFLS-R300×3, SuperBrightLEDs) at 14.6µW/mm^2^ for Throttle-gal4 and 15.9µW/mm^2^ for Tortuous-gal4 (measured at 625nm). For all GtACR experiments, blue light was presented using blue LED strips (NFLS-B300×3, SuperBrightLEDs) at 36.7µW/mm^2^ (measured at 525nm). Previously recorded behavioral responses to odor were recorded using 10% ACV (Alvarez-Salvado et al. 2018). Similar responses were measured to 1% and 10% ACV in that paper.

### Immunohistochemistry and imaging

Brains were dissected in PBS and then fixed for 14 minutes in 4% paraformaldehyde in PBS. After fixing, brains were washed 3 times in PBS and stored at 4°C until staining (≤1 week). Brains were incubated in 5% normal goat serum in PBST (0.2% Triton-X in 1x PBS) for a minimum of 60 minutes and then incubated overnight in primary antibody. Brains were then washed 3x with PBST and incubated overnight in secondary antibody. Brains were washed 3x in PBST, then mounted, covered in vectasheild (Vector Labs H-1000) and enclosed with a coverslip prior to imaging with a 20x objective (Zeiss W Plan-Apochromat 20x/1.0 DIC CG 0.17 M27 75mm) on a Zeiss LSM 800 confocal microscope at 1.25 μm depth resolution. All images were created in Image J using a maximum z-projection. Antibodies used were as follows:

| Antibodies | Source | Dilution | Identification |
| --- | --- | --- | --- |
| Mouse anti-nc82 | DSHB | 1:50 | RRID: AB_2314866 |
| Chicken anti-GFP | Fisher Scientific | 1:50 | RRID: AB_1074893 |
| Rabbit anti-DsRed | Clontech | 1:500 | RRID: AB_477652 |
| Rabbit anti-VGluT | A. DiAntonio lab | 1:1000 |  |
| Alexa488-conjugated goat anti-chicken | Fisher Scientific | 1:250 | RRID: AB_2534096 |
| Alexa633-conjugated goat anti-mouse | Fisher Scientific | 1:250 | RRID: AB_2535719 |
| Alexa568-conjugated goat anti-rabbit | Fisher Scientific | 1:250 | RRID: AB2315774 |

### Data Analysis

#### Analysis of Behavioral Data

All analysis was performed in MATLAB (Mathworks, Natick, MA).

#### Analysis of wind tunnel data

For wind tunnel experiments, individual trials were excluded due to tracking error or if flies did not move a minimum distance of 25mm over the duration of the trial. For angular velocity autocorrelogram computations, raw data (both new and from previous publications) were re-imported without low-pass filtering. For other analyses, position and orientation data were low-pass filtered at 2.5 Hz using a 2-pole Butterworth filter prior to subsequent analysis. Movement parameters including ground speed and angular velocity were computed from centroid position and orientation measurements as described previously (Alvarez-Salvado et al. 2018).

Histograms of ground speed or angular velocity were computed for each fly based on data during the first 10 seconds of baseline or post-odor periods of all trials. Data shown represent the mean and standard error across flies. Angular velocity autocorrelograms were computed from the first 10 seconds of baseline and post-odor periods of all trials. Here, raw (unfiltered) angular velocity measurements for each fly were mean-subtracted and the autocorrelogram was computed using the MATLAB function ‘xcorr’ normalized to one at zero lag. Data shown represent the mean and standard error across flies. For simulated trajectories, histograms of ground speed and angular velocity, as well as autocorrelations of angular velocity, were created using 30 seconds (1500 samples) of simulated data from each set of model parameters. Standard error was not calculated for analysis of simulated data.

To compute angular dispersion, the initial heading was subtracted from the unwrapped orientation for each time period (baseline or post-odor period) and the absolute value was taken. Data shown represent mean and standard error across flies as a function of time since the start of that period. To compute inter-turn interval distributions, turns were detected using positive threshold crossings at 45 deg/sec in the absolute angular velocity trace. Turn times were used to compute inter-turn intervals.

To segment ground speed into epochs of different means, we used a change point detection algorithm (Matlab’s ‘findchangepts’) with a minimum distance of 100 samples and a minimum threshold for residual error improvement of 1000. To analyze odor-evoked trajectories as a function of initial behavioral state, we examined the mean ground speed as determined by this algorithm in the 2 seconds before the onset of the odor stimulus. We used this value to categorize trials into ‘moving’ or ‘not moving’ using a threshold of 1mm/s.

To measure interactions between turning and groundspeed, we first detected turns as positive threshold crossings in the absolute angular velocity at 45 degrees/s, and extracted turns as the period from 0.5 seconds before to 0.7 seconds after this threshold crossings. We used the mean ground speed in the first 0.5 seconds of our extracted windows to sort both ground speed and angular velocity windows into 6 categories (<1mm/s, 1-3mm/s, 3-8mm/s, 8-12mm/s, 12-18mm/s and >18mm/s). We then calculated the mean and standard error of groundspeed and angular velocity within each category.

For the cross-correlation of model units, unit activity was mean-subtracted and the cross-correlation was calculated using Matlab’s ‘xcorr’ function using the ‘coeff’ flag for normalization.

To compute timecourses and means of locomotor parameters in Figures 4 and 5, periods when flies were moving at less than 1mm/s were omitted as done previously (Alvarez-Salvado et al. 2018). For comparisons of angular velocity at odor offset (Fig. 4E,F) we compared mean absolute angular velocity for all moving datapoints (ground speed > 1mm/s) over a 5 second period following the end of the odor period. To examine the effect of optogenetic activation or silencing on forward velocity and angular speed (Fig S4C,D, 5G,H, S5 D, S6C) we computed the mean of each parameters for moving datapoints (ground speed > 1mm/s) over the 10 second light period and compared this to the 10 second period immediately preceding the light stimulus. To compute pathlength with and without optogenetic activation or silencing (Fig. 5E) we computed the sum of forward velocity divided by the sample rate during the 10 second odor period for trials with and without light.

#### Warping and alignment of confocal images and hemibrain neurons

To determine connectome identities of neurons in split-Gal4 lines we first obtained high-resolution confocal stacks, then warped these to the JFRC2018F standard brain (Bogovic et al. 2018) using CMTK (Kohn et al. 2013, https://github.com/jefferis/fiji-cmtk-gui). We determined candidate hemibrain neurons by visual inspection using neuprint (Scheffer et al. 2020) and generated images of these neurons also warped to the JFRC2018F standard brain using the hemibrain_to_nrrd (https://github.com/wilson-lab/nat-tech) within natverse (Bates et al. 2020). We overlaid candidate neurons with our warped confocal using FIJI and determined the best fit by visual overlap.

#### Connectomics

Connectomic data from hemibrain v1.2.1 were obtained from neuprint (https://neuprint.janelia.org, hemibrain:v1.2.1) using custom queries and organized and analyzed in Microsoft Excel. We counted recurrent synapses between neurons within our identified populations, synapses from MBONs and PFL neurons onto our identified populations, and synapses from our identified populations onto DNs. Synaptic weights less than 3 were excluded.

#### Model construction and analysis

All modeling was performed in MATLAB. Our model of locomotor control was based on two layers. In the first layer, 5 units (*u*_1−5_), receive independent Gaussian noise and interact through a matrix of synaptic weights. The activity of each unit evolves according to:

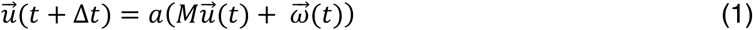

where Δ*t*= 20*ms*, equivalent to our sampling rate for behavioral data, *M* is a matrix of weights, 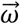 is a Gaussian noise term, and *a* is a sigmoid function that keeps activity bounded:

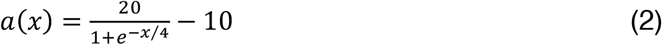

The weight matrix *M* has positive values along the diagonal, corresponding to auto-regressive terms *A*, and two negative parameters corresponding to stop inhibition (*I*_*s*_, mutual inhibition between *u*_3_ and all other units) and contralateral inhibition (*I*_*c*_, mutual inhibition between left *u*_1,2_ and right *u*_4,5_ movement-producing units):

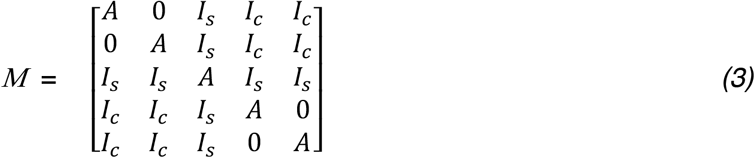

In the version of the model shown in Figure 3, *A* = 0.8, *I*_*S*_ = −0.03, *I*_*C*_ = −0.015 at baseline and *I*_*C*_ = −0.025 during offset search. As indicated in the Appendix, these values of inhibitory strength correspond to a reduction by a factor of *e* in ∼ 0.5 s for *I*_*s*_ and in ∼1 s for *I*_*c*_. Values of *a* (eq. 1) and *A* are scaled for convenience so that activity levels are in the range ±10; the Appendix analyzes model behavior after a dimensional reduction that scales activity levels to the range ±1 (which does not alter model behavior).

In the second layer, locomotor parameters (forward and angular velocity) are computed based on the activity of all units, according to the following equations:

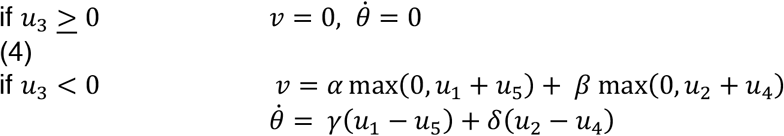

Where *ν* and 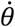 are forward and angular velocity and the parameters *α, β, γ, δ*, set the gain of forward and angular velocity for each pair of units:

|  |  |
| --- | --- |
| $\alpha$ | 1.6 mm/sample |
| $\beta$ | 0.5 mm/sample |
| $\gamma$ | 0.2 °/sample |
| $\delta$ | 0.35 °/sample |

In order to model odor effects on behavior, we introduced two odor-gated terms that impact unit activity. In the presence of odor, we added a negative offset to the stop unit to suppress its activity:

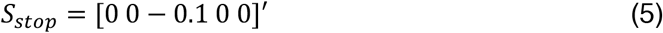

In addition, a steering function proportional to the sine of the difference between the current heading (*θ*) and the wind angle (*ψ*) was added to the activity of the movement producing units:

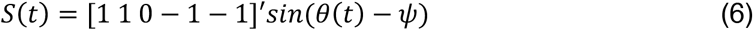

Both terms were added to the unit activity as follows:

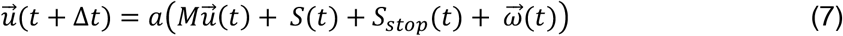

To model the effect of feedforward inhibition and activation in our model, we added a tonic offset *T*(*t*) to activity across all units:

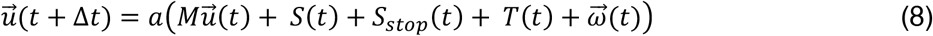

In the versions of the model testing the impact of ipsilateral connections, the weight matrix M maintains the values corresponding to both the auto-regressive terms and stop inhibition.

Contralateral inhibition is set to zero and a new term, corresponding to ipsilateral connections, replaces the zeros in the original matrix:

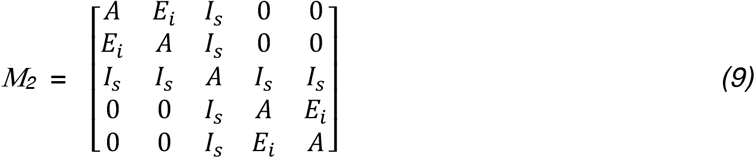

See attached appendix for detailed analysis of the model.

## Key Resources Table

| Resource | Source | Identifier |
| --- | --- | --- |
| **FLY LINES** |  |  |
| VGlut-AD | BDSC | RRID:BDSC_84713 |
| VT043158-DBD | BDSC | RRID:BDSC_73583 |
| 31A11-DBD | BDSC | RRID:BDSC_69813 |
| SS49952 | BDSC | RRID:BDSC_88573 |
| SS41947 | BDSC | RRID:BDSC_86951 |
| SS38176 | BDSC | RRID:BDSC_87082 |
| VT047747 AD; VT043158DB | This study | RRID:BDSC_601020<br>RRID:BDSC_73583 |
| UAS-Chrimson | BDSC | RRID:BDSC_55136 |
| w1118; s/Cyo; UAS-GtACR1-EYFP/TM6b | Claude Desplan |  |
| UAS-GtACR1/CyO | BDSC | RRID:BDSC_92988 |
| 10X UAS-CD8:GFP | BDSC | RRID:BDSC_32183 |
| **REAGENTS** |  |  |
| All trans retinal | Sigma | RRID: R2500 |
| **SOFTWARE** |  |  |
| Python |  | Version 3.8 |
| MATLAB | Mathworks | Version R2023a |
| Image J |  | Version 1.53t |
| Microsoft Excel |  | Version 2308 |
| FicTrac |  | Version 2.0 (build date: Nov 2 2018) |
| CMTK Registration | https://github.com/jefferis/fiji-cmtk-gui |  |
| R/Rstudio | https://cran.rstudio.com<br>https://posit.co/download/rstudio-desktop/ |  |
| natverse | https://natverse.org |  |
| hemibrain_to_nrrd | https://github.com/wilson-lab/nat-tech |  |
| **DATASETS** |  |  |
| hemibrain connectome | https://neuprint.janelia.org/ | Version: v1.2.1 |
| Data from Alvarez-Salvado et al. 2018 | https://doi.org/10.5061/dryad.g27mq71 |  |

---

Supplementary Information

**Supp Fig 1:**
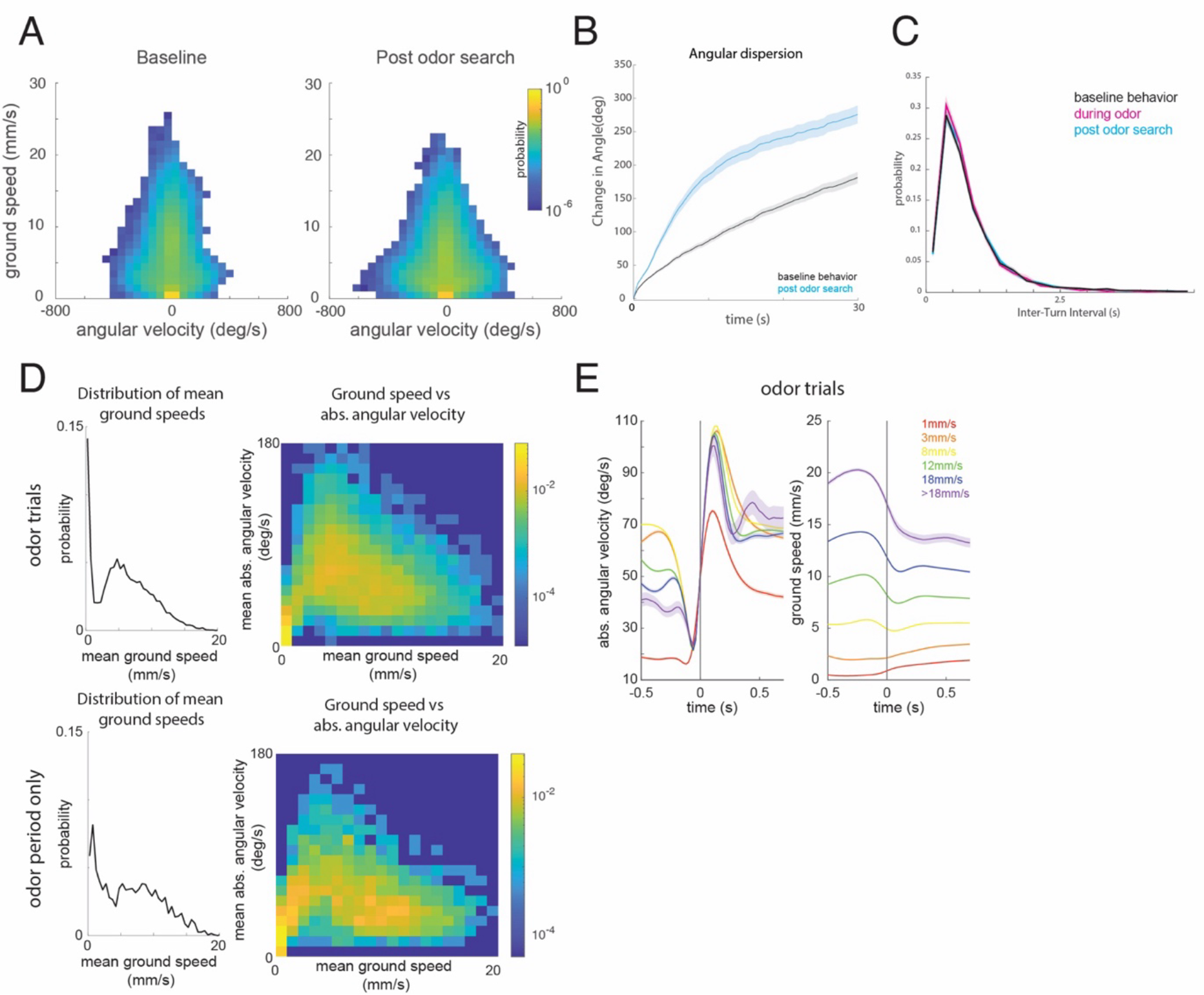
A. Two dimensional histograms of ground speed and angular velocity for the first 10 seconds of baseline behavior (left) and the first 10 seconds of post-odor search (right). Plots represent the mean histogram across flies (n=1306 total trajectories from 75 flies). Probability on the color axis. Same flies as in Fig 1D-F. Flies favor slower ground speeds and larger angular velocities after odor loss. B. Angular dispersion as a function of time relative to heading at the start of trial (black, baseline period) or odor loss (cyan: post-odor period). Traces represent mean ± standard error across flies (n=1306 total trajectories from 75 flies). Same flies as in Fig 1D-F. Flies turn further after loss of odor when compared to baseline behavior. C. Distribution of inter-turn intervals during baseline (black), odor (magenta), and post-odor periods (cyan) periods. Error bars represent standard error across flies (n=1306 total trajectories from 75 flies). Same flies as in Fig 1D-F. D. Distribution of mean ground speeds and joint distribution of mean ground speed and mean angular velocity for odor trials (top) and odor periods only (bottom). The relationship between groundspeed and angular velocity is similar to those observed in unstimulated trials (Fig. 2C,D) except that fewer flies are stationary during odor periods. E. Angular velocity (left) and ground speed (right) relative to turn onset for odor trials as in Fig. 2E (n=1306 trajectories from 75 flies).

**Supp Fig 2:**
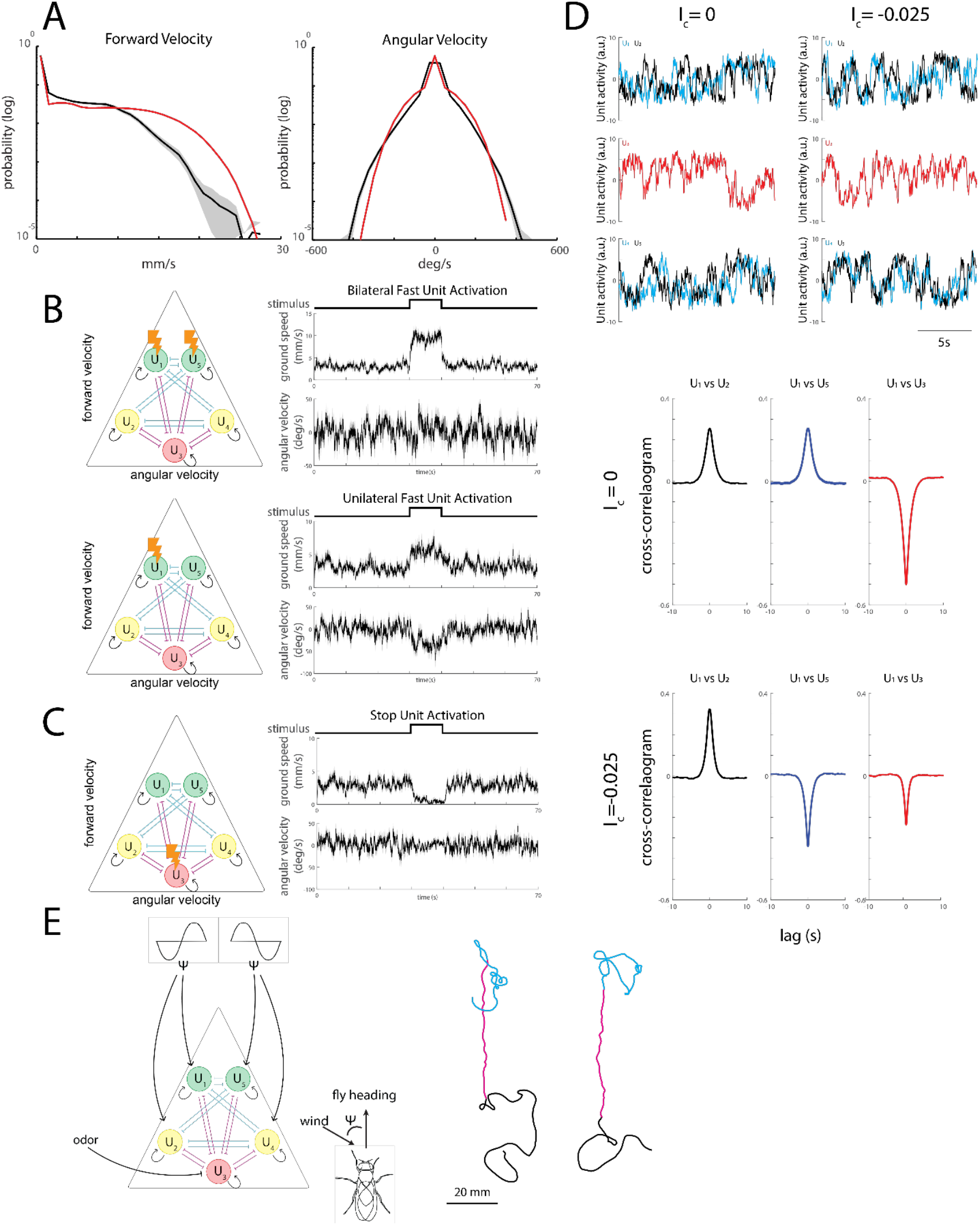
A. Distributions of ground speed (left) and angular velocity (right) produced by the model (red) after tuning the model coefficients (α,β, γ, δ), overlaid on experimental distributions (black) for baseline walking. Model data computed from 1300 generated trajectories. Baseline data reproduced from Fig.1D,E. B. Effects of unit activation on locomotor parameters. Top: bilateral activation of *u*_1_ and *u*_5_ drives an increase in ground speed with no net change in angular velocity. Bottom: unilateral activation of *u*_1_ drives turning, indicated here by a negative shift in angular velocity, and a smaller increase in ground speed. C. Effects of stop unit activation on locomotor parameters. Activation of the stop unit *u*_)_ drives an abrupt decrease in ground speed. D. Unit activity in the absence (left) and presence (right) of contralateral inhibition (*I*_*c*_). In the absence of contralateral inhibition activity in a locomotor unit (*u*_1_) is correlated both with the unit on the same side (*u*_2_) and on the opposite side (*u*_5_), but is negatively correlated with activity in the stop unit (*u*_3_)— quantified in middle plot below. In the presence of contralateral inhibition (right) activity in units on opposite sides becomes anti-correlated – quantified in bottom plot – resulting in larger angular velocities and smaller forward velocities. E. Model schematic with the addition of odor- and wind-gated terms (left). Odor suppresses the activity of the stop-producing unit (*u*_3_). Heading is compared to wind direction to generate upwind turn commands in movement-producing units (*u*_1_, *u*_2_, *u*_4_, *u*_5_). Resulting trajectories (right). Black: baseline walking, magenta: odor, cyan: post-odor search.

**Supp Fig 3:**
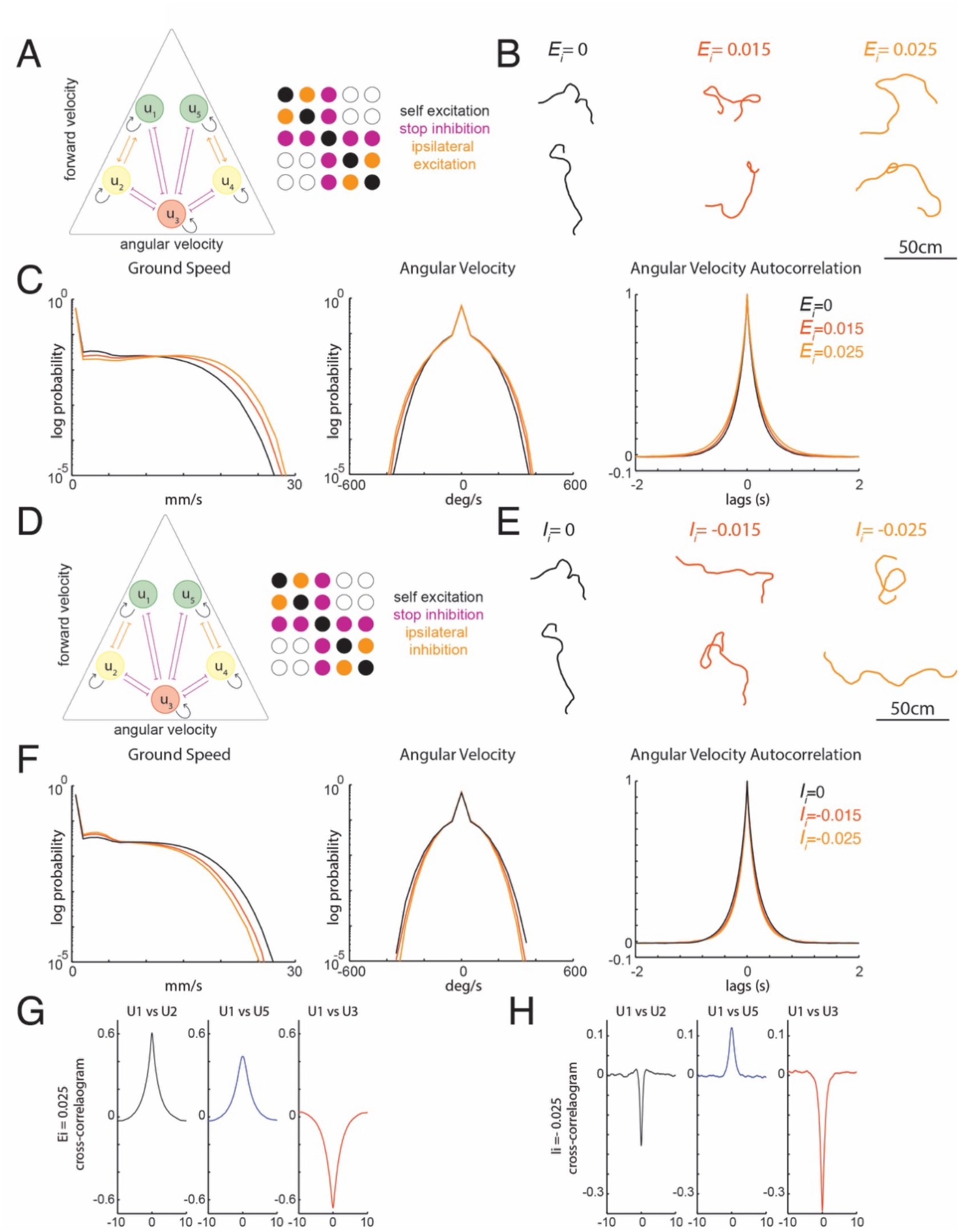
A. Model variant with ipsilateral excitation instead of contralateral inhibition. Symbols as in Fig. 3A-C. B. Example trajectories of model flies with different amounts of ipsilateral excitation (*E*_*i*_). C. Statistical features of locomotion in model flies (from 1300 model trajectories) as a function of the strength of ipsilateral excitation. Left: Ground speed distribution, center: angular velocity distribution, right: autocorrelation of angular velocity. Increased ipsilateral excitation shifts both the ground speed and angular velocity distributions towards higher values and slightly widens the autocorrelogram of angular velocity. D. Model variant with ipsilateral inhibition. E. Example trajectories of model flies with different amounts of ipsilateral inhibition (*I*_*i*_). F. Statistical features of locomotion in model flies (from 1300 model trajectories) as a function of the strength of ipsilateral inhibition. Left: Ground speed distribution, center: angular velocity distribution, right: autocorrelation of angular velocity. Increased ipsilateral inhibition shifts both the ground speed and angular velocity distributions towards lower values and slightly narrows the autocorrelogram of angular velocity. G. Quantification of the correlation between one unit (*u*_1_) and a unit on the same side of the model (*u*_2_) (left), opposite side of the model (*u*_5_) (middle) and the stop unit (*u*_3_) (right) with the addition of ipsilateral excitation. Ipsilateral excitation drives correlation between activity across all movement producing units. Activity in *u*_1_ is negatively correlated with the stop unit. H. Quantification of the correlation between one unit (*u*_1_) and a unit on the same side of the model (*u*_2_) (left), opposite side of the model (*u*_5_) (middle) and the stop unit (*u*_3_) (right) with the addition of ipsilateral inhibition. Ipsilateral inhibition produces anticorrelated activity with units on the same side of the model and correlated activity with units on the other side. Activity in *u*_1_ is negatively correlated with the stop unit.

**Supp Fig 4:**
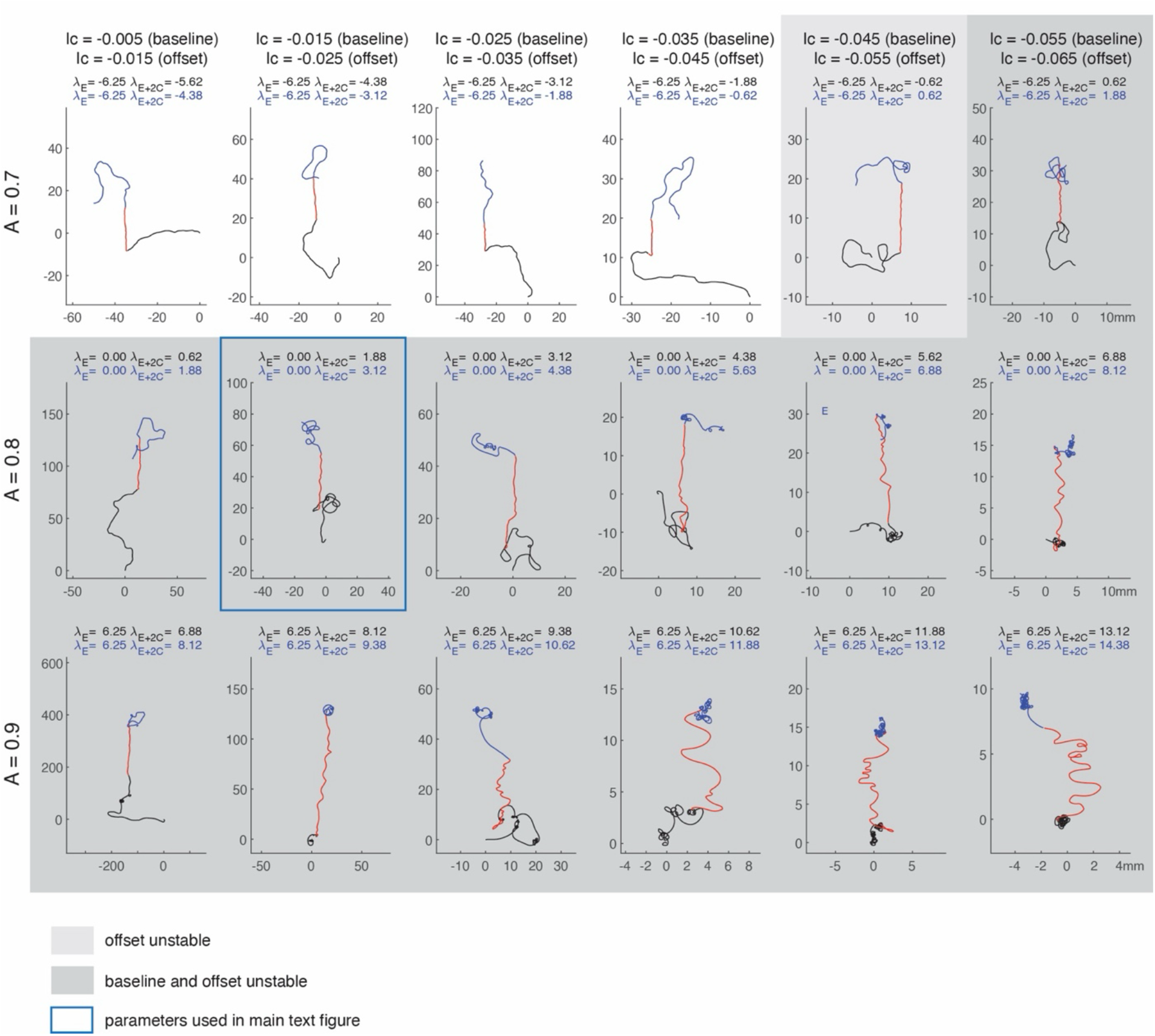
Trajectories produced by the model in the main text, as the self-excitation parameter A is varied (top to bottom: 0.7, 0.8, 0.9) and the contralateral inhibition parameter *I*_*s*_ is varied (left to right: - 0.005 to -0.055 in steps of -0.01). As in the main text, *I*_*s*_ = −0.03, *I*_*c*_ is made more negative by 0.01 after odor offset, and the odor is off for 30 sec (black), then on for 10 sec (red), then off 30 sec (blue). Other model parameters are unchanged. A gray background indicates that one or both phases of the trajectory have an unstable anti-symmetric component, as shown in the legend. The second panel in the second row corresponds to the model parameters in the main text (blue rectangle). Numbers in each panel indicate the eigenvalues for the fixed point of the antisymmetric component for baseline (black) and odor-offset (blue) parameters; positive values are unstable. Note the change in overall trajectory size, as indicated by the axis tick-marks.

**Supp Fig 5:**
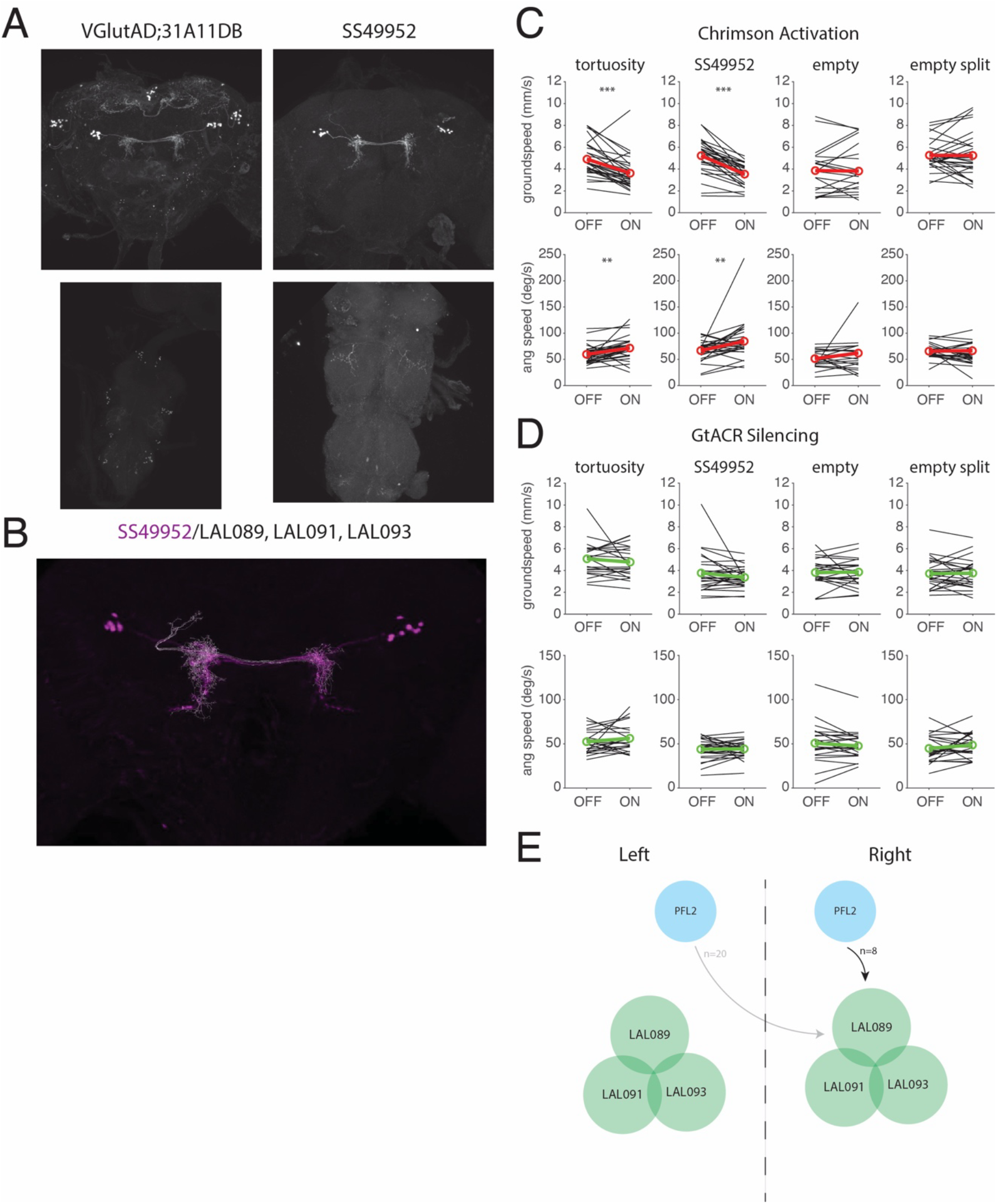
A. Confocal images of neurons labeled by VGlutAD;31A11DB > GFP (green) and SS49952 > GFP (green) in the brain (top) and ventral nerve cord (bottom). GFP signal only. B. Warped confocal image of neurons labeled by SS49952 > GFP (magenta) overlaid with best-fit neurons from the hemibrain connectome: LAL089 (n=2/side), LAL091 (n=4/side), and LAL093 (n=2/side). C. Mean groundspeed and absolute angular velocity of each line analyzed in Figure 4 expressing Chrimson in the absence (OFF) versus presence (ON) of light. Black lines: average for each fly. Red lines: average across flies. Asterisks: statistically significant changes in paired *t*-tests. VGlutAD;31A11DB (tortuosity) > Chrimson (g.s. p = 2.9162E-05, ang vel p = 0.0042683, 183 total trials from 31 flies), SS49952 > Chrimson (g.s. p = 4.2227E-09, ang vel p = 9.2759E-03, 250 total trials from 31 flies), empty-Gal4 > Chrimson (g.s. p = 0.51294, ang vel p = .47303, 411 total trials from 25 flies) and empty Split-Gal4 > Chrimson (g.s. p = 0.94536, ang vel p = 0.18964,146 total trials from 30 flies). Groundspeed data for empty-Gal4 and empty-Split Gal4 are also shown in Figure 5. D. Mean groundspeed and absolute angular velocity of each line analyzed in Figure 4 expressing GtACR in the absence (OFF) versus presence (ON) of light. Black lines: average for each fly. Green lines: average across flies. Asterisks: statistically significant changes in paired *t*-tests. VGlutAD;31A11DB (tortuosity) > GtACR (g.s. p = 0.31282, ang vel p = 0.2546, 199 total trials from 23 flies), SS49952 > GtACR (g.s. p = 0.18082, ang vel p = 0.70713, 279 total trials from 29 flies), empty > GtACR (g.s. p = 0.69946, ang vel p = 0.22707, 224 total trials from 29 flies), and empty split > GtACR (g.s. p = 0.97136, ang vel p = 0.1748, 227 total trials from 30 flies) in the absence (OFF) and presence (ON) of light. Groundspeed data for empty-Gal4 and empty-Split Gal4 are also shown in Figure 5. E. Connectomic analysis of neurons labeled by SS49952. These neurons form both input and output arbors in the medial LAL without notable projections outside of this zone. This population receives mild bilateral innervation from PFL2 neurons in the Central Complex.

**Supp Fig 6:**
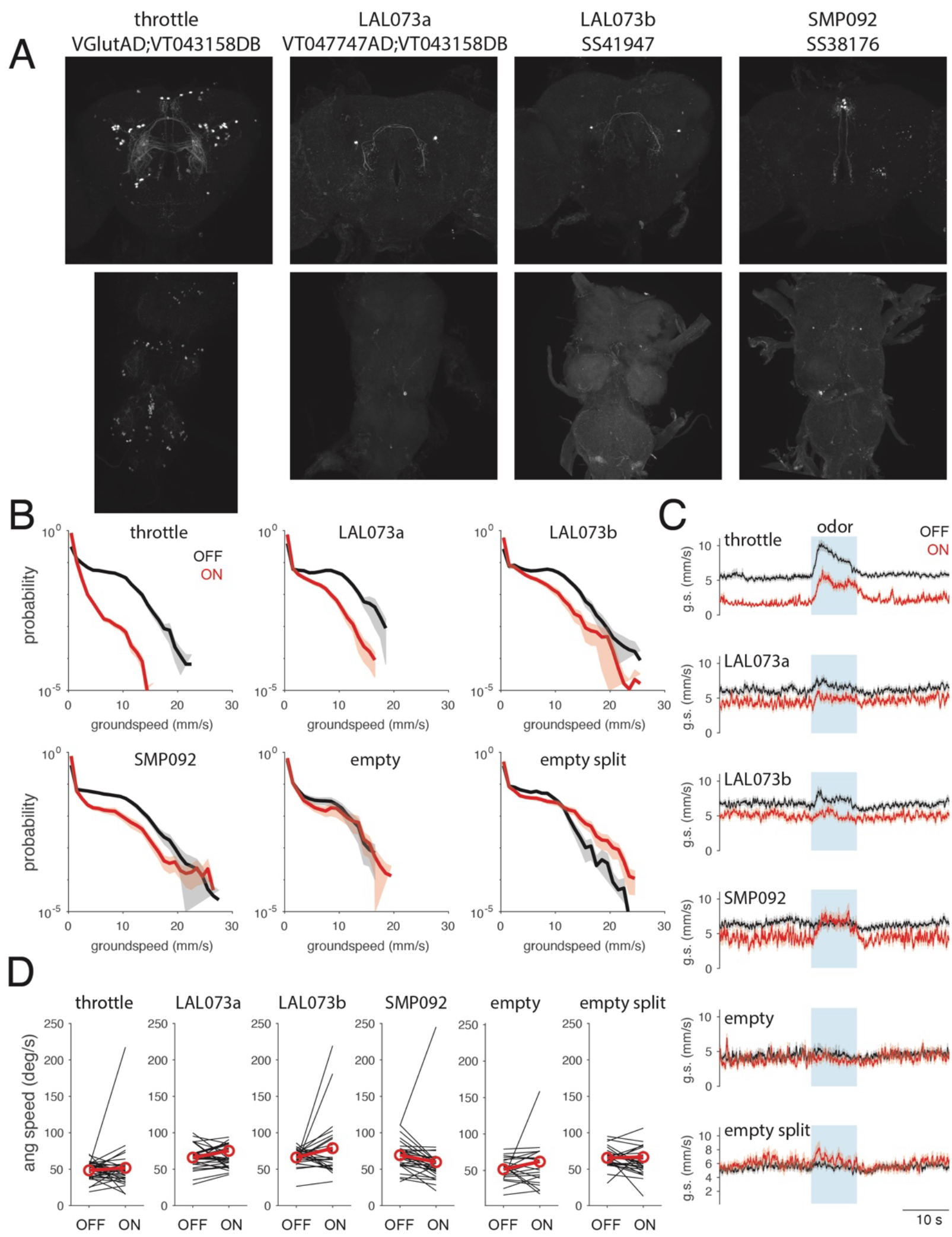
A. Confocal images of VGlutAD;VT043158DB (throttle) > GFP, VT047742;VT043158DB > GFP, SS41947 > GFP, and SS38176 > GFP neurons labeled in the brain (top) and ventral nerve cord (bottom). GFP signal only. B. Groundspeed distribution in the absence (black) and presence (red) of light for each line expressing Chrimson. Error bars represent standard error across flies. Activation shifts groundspeeds towards lower values for VGlutAD;VT043158DB (throttle), VT047742;VT043158DB (LAL073a), SS41947 (LAL073b), and SS38176 (SMP092), but not for empty-Gal4 or empty split-Gal4. Numbers of trials and flies as in Fig. 5G. C.Time course of mean groundspeed (+/- S.E.) for each line expressing Chrimson and responding to an odor pulse (blue) in the absence (black) and presence (red) of light. Note the strong reduction in mean groundspeed in the VGlutAD;VT043158DB (throttle) line, and weaker reduction in groundspeed in VT047742;VT043158DB (LAL073a), SS41947 (LAL073b), and SS38176 (SMP092). Numbers of trials and flies as in Fig. 5G. D. Mean absolute angular velocity for each line analyzed in Figure 5 expressing Chrimson in the absence (OFF) versus presence (ON) of light. Black lines: average for each fly. Red lines: average across flies. Activation produced no significant change in absolute angular velocity in any of these lines (p = 0.55803, 0.90872, 0.15721, 0.11388, 0.47303, 0.18964 respectively). Numbers of trials and flies as in Fig. 5G.

**Supp Fig 7:**
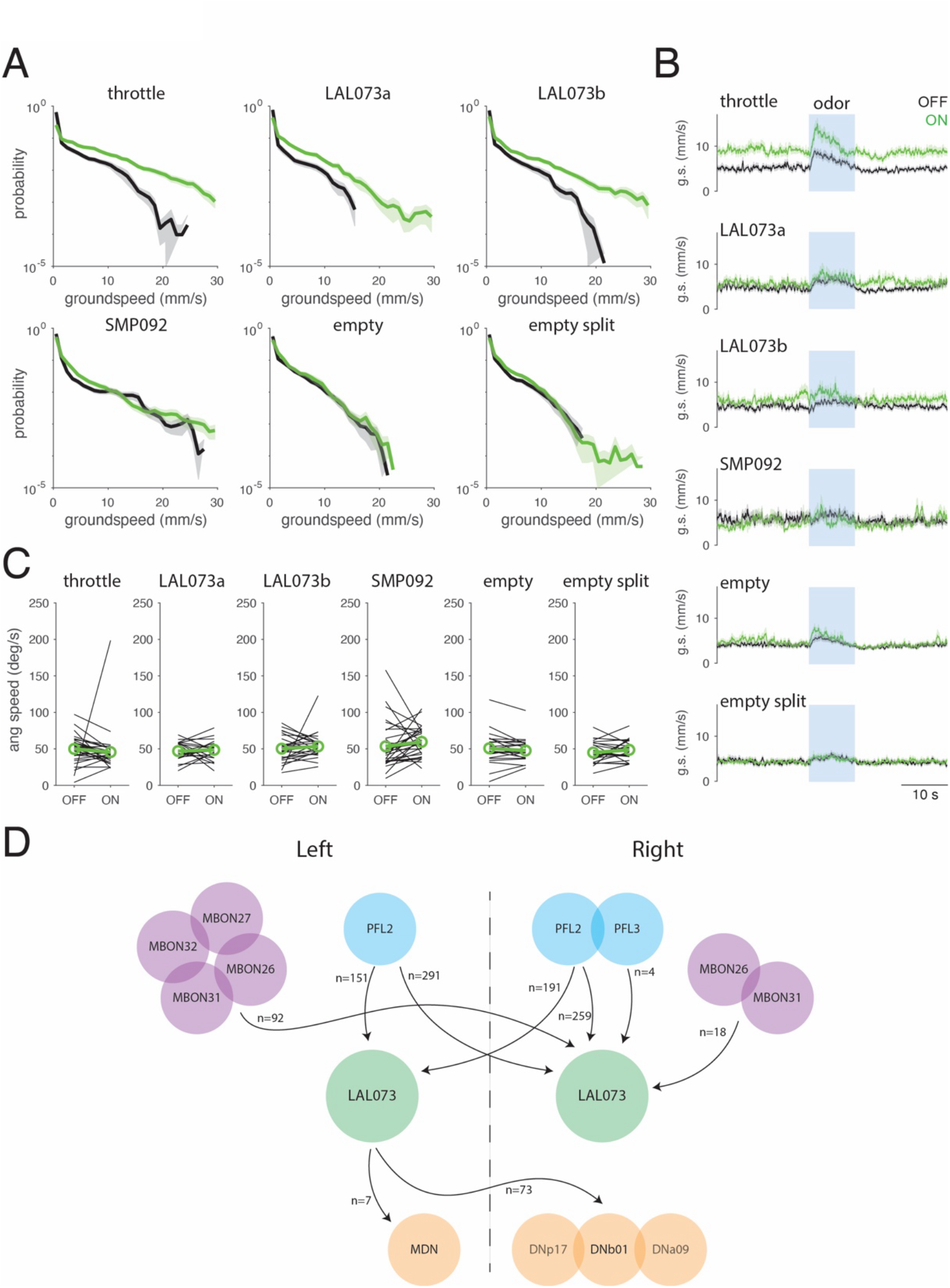
A. Groundspeed distribution in the absence (black) and presence (green) of light for each line expressing GtACR. Error bars represent standard error across flies. Activation shifts groundspeeds towards higher values for VGlutAD;VT043158DB (throttle), VT047742;VT043158DB (LAL073a), and SS41947 (LAL073b), but not for SS38176 (SMP092), empty-Gal4, or empty split-Gal4. Numbers of trials and flies as in Fig. 5H. B. Time course of mean groundspeed (+/- S.E.) for each line expressing GtACR and responding to an odor pulse (blue) in the absence (black) and presence (green) of light. Note the strong increase in mean groundspeed in the VGlutAD;VT043158DB (throttle) line, and weaker increase in groundspeed in the VT047742;VT043158DB (LAL073a) and SS41947 (LAL073b) lines. Numbers of trials and flies as in Fig. 5H. C. Mean absolute angular velocity for each line analyzed in Figure 5 expressing GtACR in the absence (OFF) versus presence (ON) of light. Black lines: average for each fly. Green lines: average across flies. Activation produced no significant change in absolute angular velocity in any of these lines (p = 0.64588, 0.84898, 0.35746, 0.29879, 0.22707, 0.1748 respectively). Numbers of trials and flies as in Fig. 5H. D. Connectomic analysis of LAL073. This neuron receives very strong bilateral input from PFL2 neurons, and moderate bilateral input from select MBONs, with weak input from PFL3. It makes outputs to DNs MDN, DNp17, DNb01, and DNa09.

**Table 1:**
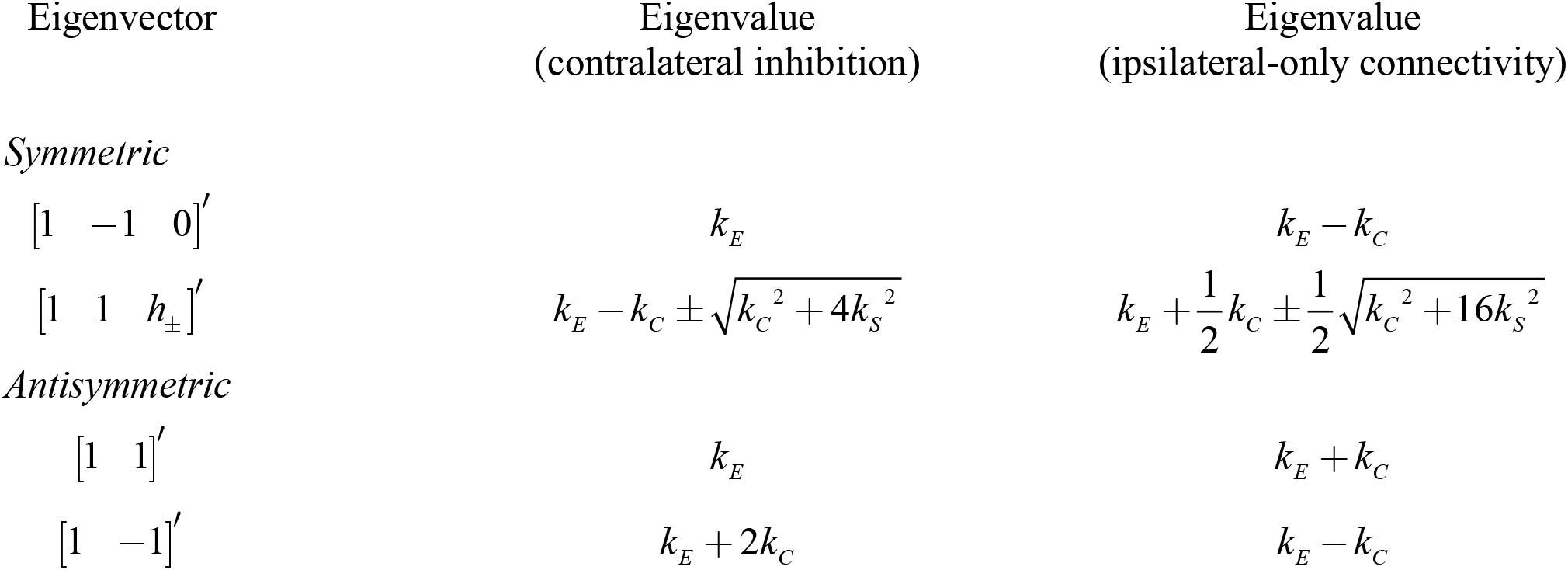
eigenvectors and eigenvalues of two models.

## Appendix Locomotion Control Model

This appendix provides an approximate analytic treatment of the model of locomotion control presented in the body of the manuscript, with the goal of providing insights into the model’s behavior and its dependence on parameters and connectivity.

The primary result is that the model can be broken into two components (“symmetric” and “antisymmetric”) that separately control forward velocity and angular velocity, and are differentially impacted by the control signals related to odor and wind that we introduce. The dynamic characteristics of the antisymmetric component are directly manifest in the tortuosity of the trajectories produced by the model and are controlled by the eigenvalues of that component, with a positive eigenvalue leading to unstable behavior and more tortuous trajectories. We find that when self-excitation is set close to the border between stable and unstable behavior, the level of contralateral inhibition can more readily and smoothly impact tortuosity – an observation that motivates our choice of parameters in the main text. We then show that the model exhibits similar dynamics regardless of the number of locomotor units on each side, so long as the symmetries of the connectivity matrix are preserved. Finally, we show how the behavior of this model contrasts with that of an alternative model, in which contralateral inhibition is replaced by ipsilateral connectivity (either excitatory or inhibitory).

A more detailed outline of the analysis of the model in the main text is as follows. We begin by showing that for small signals, the model can be approximated by a linear differential equation for a five-dimensional vector of unit activities. The five-dimensional model splits into a three-dimensional symmetric part that is concerned with forward velocity and a two-dimensional antisymmetric part that is concerned with angular velocity, and this splitting is exact for the linearized approximation. As they are linear, both of these components have a single fixed point. Depending on parameter values, the fixed points of the two subsystems can be stable, neutrally stable, or unstable, and we determine the parameter values at which these transitions occur. (In the unstable regime, activity is nevertheless bounded, because of the model’s sigmoidal activation function). For the antisymmetric subsystem, the stable regime produces trajectories are straighter than what is observed experimentally. Parameters that lead to a slightly unstable fixed point provide good matches for experimental data by producing periods of consistent turning in one direction or the other. Increasing the strength of self-excitation or contralateral inhibition pushes the fixed point of the antisymmetric component further into instability, further elongating the periods of consistent turning to a duration that results in trajectories that are too curved to match experimental data. While increasing the strength of self-excitation or stop-inhibition can also lead to instability, it does not create the observed turning behavior, and the analysis shows why this is the case: these destabilizing influences primarily affect the symmetric subsystem, which controls forward but not angular velocity. We then show that the model’s behavior is preserved with either more, or fewer, units on each side. Specifically, changing the number of units has the same effect on stability characteristics as changing the per-connection strength of the inhibitory signals.

### Linearization and conversion from difference equation to differential equation

The model equation of the main text can be rewritten as

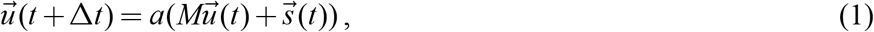

where 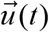 is the column vector [*u*_1_(*t*) *u*_2_ (*t*) *u*_3_ (*t*) *u*_4_ (*t*) *u*_5_ (*t*)]′, *M* is a 5×5 matrix of coupling constants (see Fig. 3C and text equation 3), 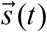 is a driving signal, and *a*(*x*) is a sigmoidal activation function to be applied element-by-element to its column-vector argument. Reproducing eq. (3) of the main text,

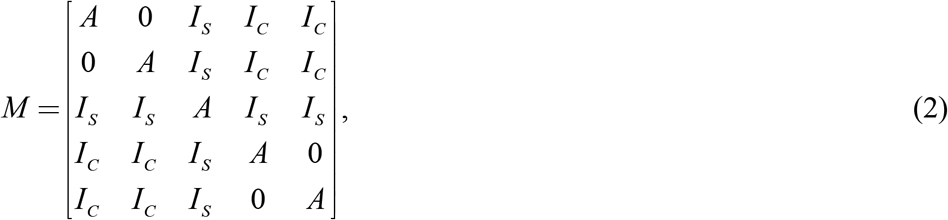

where *A* = 0.8, *I*_*S*_ = −0.03, and *I*_*C*_ = −0.015 at baseline and *I*_*C*_ = −0.025 at odor offset. The driving signal 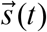 consists of independent Gaussian noise (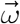 in the main text), to which exogenous signals due to odor, wind, and activation (text equations 6, 7, and 9) may be added. The activation function is specified by text equation (2), and may be rewritten as

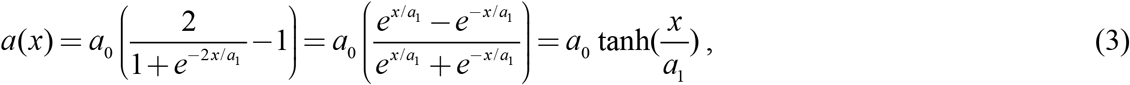

with *a*_0_ = 10 and *a*_1_ = 8. Note that when its inputs are small, the activation function is approximately linear:

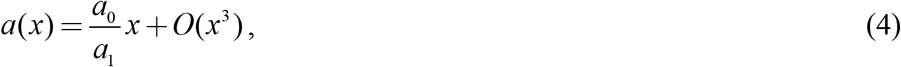

an approximation that is useful when |*x*| ≪ *a*_1_.

The model equation (1) is equivalent to

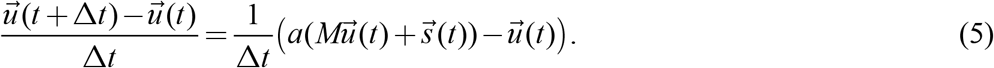

Assuming that Δ*t* is small compared to the time course of input fluctuations (Δ*t* = 20 ms in the implementation of eq. (1)), and that the inputs are sufficiently small so that eq. (4) can be used, eq. (5) is approximated by

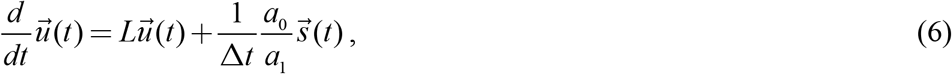

where

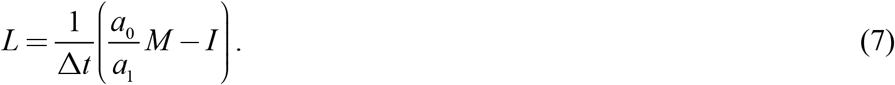

Thus, *L* is a 5×5 matrix with the same symmetry as *M*,

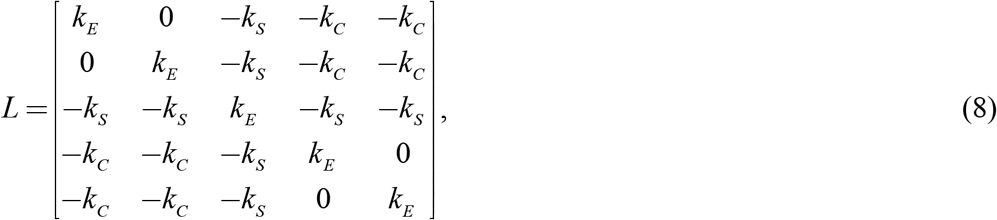

where the constants *k*_*E*_ (self-excitation), *k*_*S*_ (stop-inhibition), and *k*_*C*_ (contralateral inhibition) are related to the quantities in the main text by

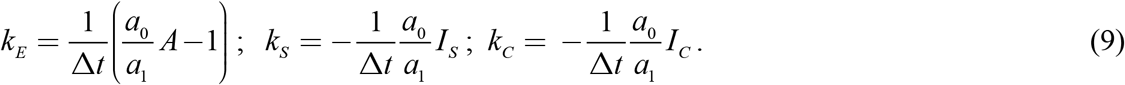

Importantly, for the parameter values considered in the main text (*A* = 0.8, *a*_0_ = 10 and *a*_1_ = 8), eq. (9) shows that the self-excitation strength *k*_*E*_ = 0. That is, the level of recurrent excitation postulated in the difference-equation formulation of the main text does not require an explicit recurrent connection at all, and corresponds to units which, if unperturbed, maintain their activity level over time.

Note also that the constants *k*_*S*_ and *k*_*C*_ are positive for the range of values of parameters considered in the main text; positive values indicate inhibition because of the minus sign in eq. (8). Eq. (9) converts these values into time rate constants: specifically, *I*_*C*_ = −0.015 corresponds to *k*_*C*_ = 0.9375 s^-1^ (i.e., a reduction by a factor of *e* in slightly more than 1 s), *I*_*S*_ = −0.03 corresponds to *k*_*C*_ = 1.87 s^-1^, a twofold more rapid and reduction.

As in Fig. 3B of the main text, the activities *u* result in locomotion whose forward and angular velocities are given by

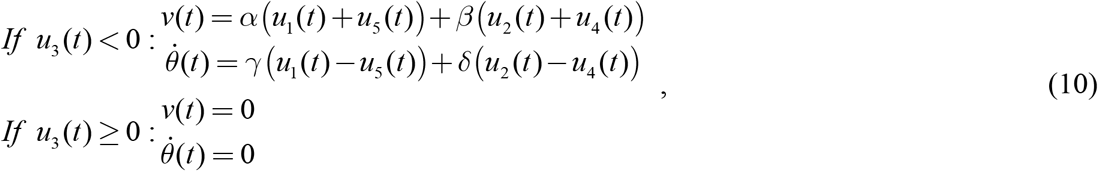

where *α, β, γ*, and *δ* are specified in the text.

#### Splitting the system

The full model is five-dimensional, but the symmetry of the coupling matrix *L* (eq. (8)) allows for a splitting of the model via a coordinate rotation:

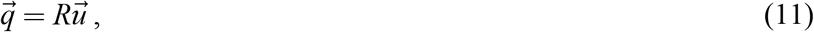

where

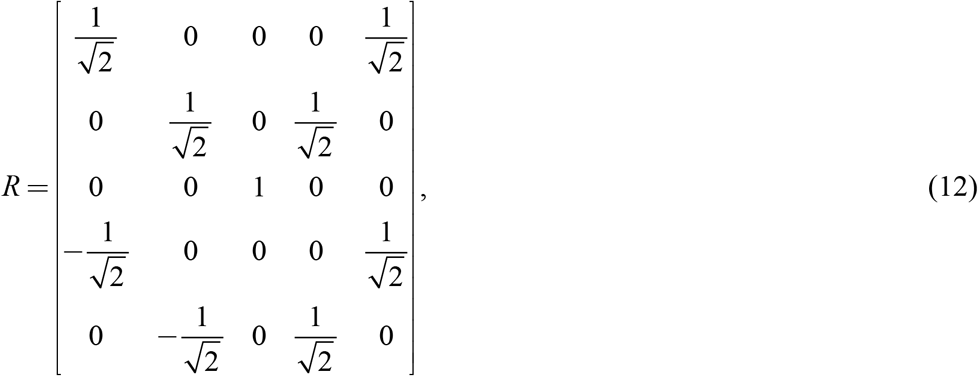

is a rotation matrix, satisfying *R*^−1^ = *R*^*T*^. In the new coordinates, the model (eq. (6)) becomes

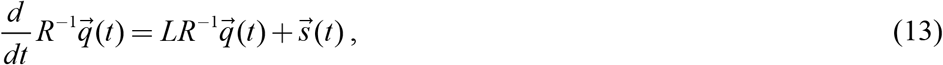

which is equivalent to

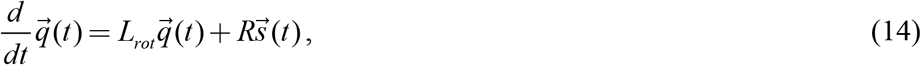

with *L*_*rot*_ = *RLR*^−1^. As expected from the symmetry-based transformation of model variables, the transformed coupling matrix *L*_*rot*_ is block-diagonal:

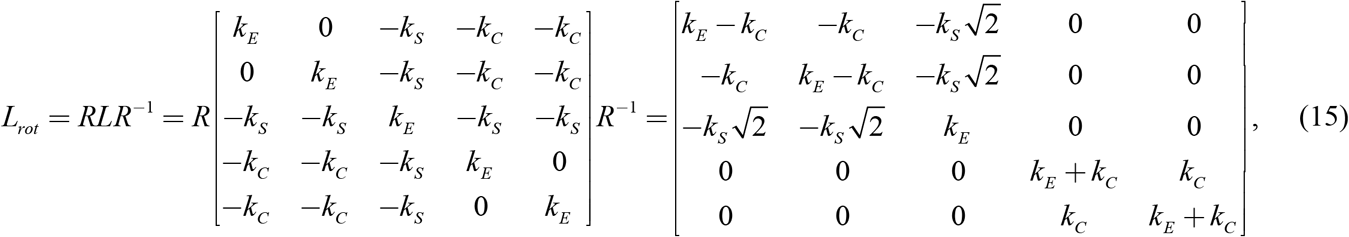

Eq. (15) shows that the transformation of the original model (eq. (6)) into rotated variables (eq. (14)) splits the original system into a three-dimensional part in which the original variables are combined symmetrically, and a two-dimensional part in which they combine antisymmetrically. We make this split explicit, with 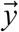 the three-dimensional symmetric part, and 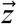 the two-dimensional antisymmetric part:

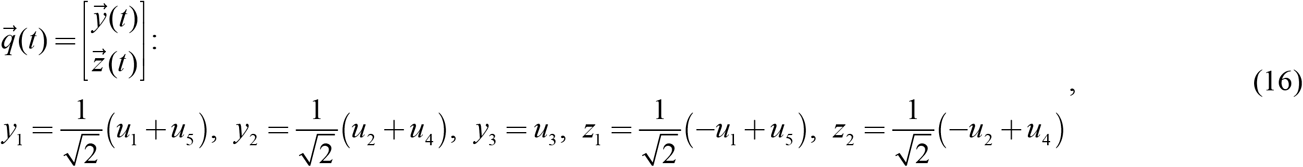

The original system is now approximated by

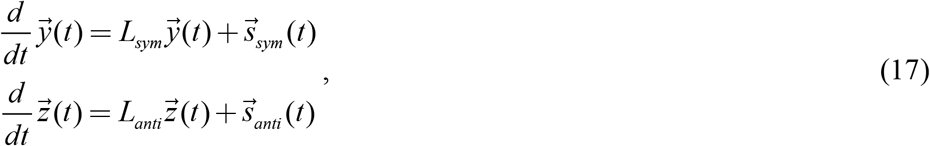

where

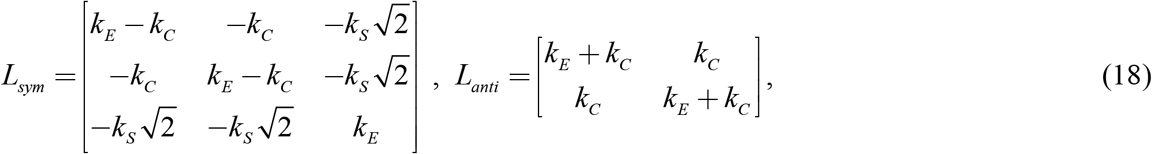

and 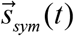 and 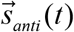 are the first three, and the last two, dimensions of 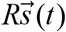.

This splitting simplifies the analysis of the model’s driving signals and locomotor outputs. Since the transformation *R* is a rotation, the noise component of 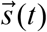, which is assumed to be a set of uncorrelated Gaussian noises, remains uncorrelated and Gaussian when split into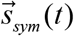 and 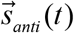. The other model inputs selectively activate either the symmetric or antisymmetric components. The stop signal, *c*_*stop*_ [0 0 −1 0 0]′ (main text eq. 5) splits into a symmetric component *c*_*stop*_ [0 0 −1]′ and an antisymmetric component of 0 (here, []′ indicates transpose.). The wind-related signal, sin(*θ*(*t*) − *ψ*)[1 1 0 −1 −1]′ (main text eq. 6) splits into a symmetric component of 0 and an asymmetric component of 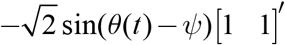. Tonic activity *c*_*tonic*_ (*t*)[1 1 1 1 1]′ splits into a symmetric component 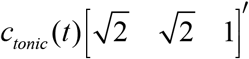 and an antisymmetric component of 0. In terms of the new variables of eq. (17), the locomotor outputs (10) become

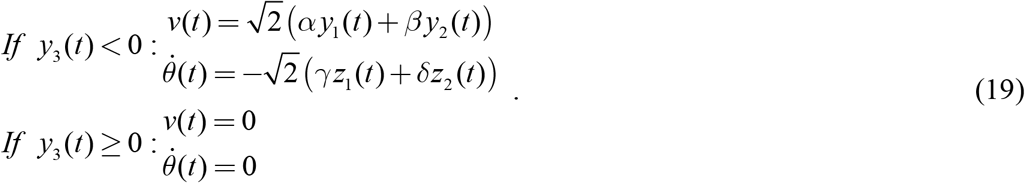

That is, the forward velocity signal only depends on the output of the symmetric component, and the angular velocity signal only depends on the output of the antisymmetric component. In the linearized approximation, the only coupling of these signals occurs when they are translated into movement: the stop signal (which is part of the symmetric component) vetoes both forward velocity and angular velocity, and the “gating” equation (main text eq. 5) by which angular velocity modulates forward velocity.

Finally, note that the symmetry-based splitting remains useful even when the small-signal linearizing approximation tanh *x* ≈ *x* used in eq. (4) does not hold, since the sigmoidal compression respects this symmetry: tanh(−*x*) =−tanh(*x*). Consequently, a model state in which activity confined to the symmetric component will remain confined to that component and generate no activity in the antisymmetric component, and vice-versa.

#### Fixed-point behavior

As is standard, we analyze the qualitative behavior of the systems given by eq. (17) in terms of the presence of fixed points and the behavior of model dynamics at the fixed points. This analysis, which assumes that the noise input is infinitesimal and uncorrelated, is expected to capture the qualitative behavior of the system under circumstances in which the excursions due to the noise are small over periods corresponding to the time constants of the system. Beyond this regime, the influences of the saturating nonlinearity (eq. (3)) and the correlations structure of the noise need to be taken into account.

Since each of these systems is a linear differential equation with constant coefficients, they each have a single fixed point, at the origin. Moreover, since the matrices in eq. (18) are real and symmetric (and non-singular), their full set of solutions are sums of exponentials 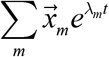, where the constants *λ* are the eigenvalues of the matrices *L*_*sym*_ and *L*_*anti*_, and *x* is a two- or three-dimensional column eigenvector. These eigenvalues determine the qualitative behavior of the equations’ solutions, and the system’s responses to pulse inputs. If all eigenvalues are negative, all solutions tend to zero and the fixed point is stable; if any *λ*_*m*_ is positive, then there are solutions 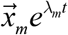 that will depart from the fixed point exponentially. In the full model, this exponential departure will continue until the linearized approximation is no longer valid, at which point the saturation of the sigmoidal activation function will keep the solution bounded.

The eigenvectors and eigenvalues for both subsystems are readily determined (below). In both cases, since the matrices *L*_*sym*_ and *L*_*anti*_ are real and symmetric, their eigenvalues must be real and their eigenvectors must be orthogonal.

#### Symmetric component

For *L*_*sym*_, [1 −1 0]′ is evidently an eigenvector, with eigenvalue *λ*_*E*_ = *k*_*E*_. The other eigenvectors, since they must be orthogonal to [1 −1 0]′, are of the form 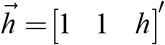.

To determine *h*, note that if 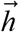 is an eigenvector, then 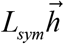 must be proportional to 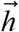, so the third element of

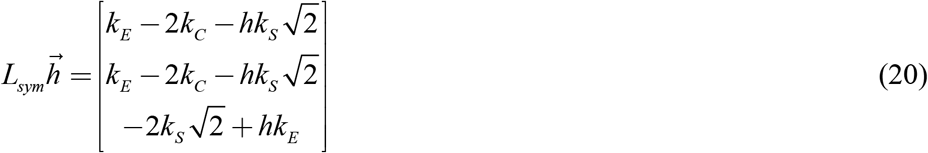

must be *h* times as large as each of the first two elements. This requires

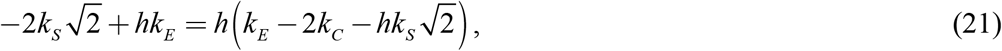

a quadratic equation whose roots are given by

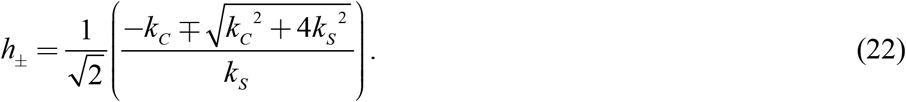

With these choices, 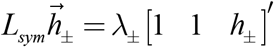. So the two remaining eigenvalues *λ*_±?_can be read off from the first entry of 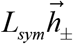 on the right-hand side of eq. (20):

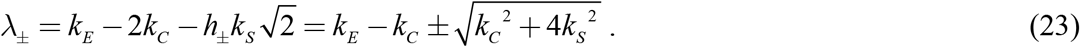

Thus, the symmetric component has one eigenvalue at *k*_*E*_, and two eigenvalues equally-spaced above and below *k*_*E*_ − *k*_*C*_, with the more positive value (*λ*_+_ in eq. (23)) always greater than *k*_*E*_. So if *k*_*E*_ > 0, at least two eigenvectors determine unstable trajectories (*λ*_*E*_ = *k*_*E*_ and *λ*_+_ in eq. (23)). Conversely, *k*_*E*_ < 0 is required for stability. However, even if *k*_*E*_ < 0, there is always a larger eigenvalue, *λ*_+_, and it too must be negative to ensure stability. That is, the symmetric component’s fixed point is only stable when *k*_*E*_ < 0 and *λ*_+_ < 0, which in turn requires that

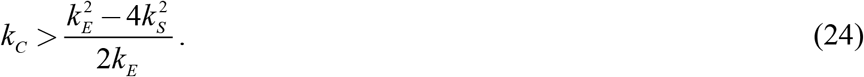

In terms of the parameters in the main text, *k*_*E*_ = 0 when 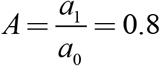 (per eq. (9)), so the condition that *k*_*E*_ < 0 equivalent to is 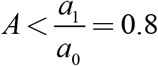. The condition (24), in terms of the model parameters of the main text, is

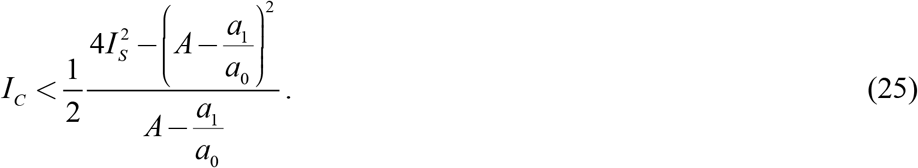

#### Antisymmetric component

The antisymmetric component determines the model’s turning behavior, and it has a simpler structure than the symmetric component. The form of the 2×2 matrix with eigenvalue *L*_*anti*_ makes both eigenvectors evident: [1 1]′, with eigenvalue

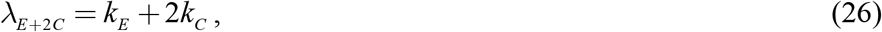

and [1−1]′ with eigenvalue *λ*_*E*_ = *k*_*E*_. Both eigenvectors result in consistent turning in a single direction, but the [1 1]′ eigenvector will result in a greater angular velocity since the two terms that contribute to 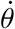 in eq. (19) will reinforce.

When *k*_*E*_ is negative (i.e.,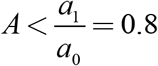) and *k*_*E*_ + 2*k*_*C*_ < 0, the sole fixed point at the origin is stable.

Thus, in terms of the model parameters of the main text, the fixed point of the antisymmetric component is stable when

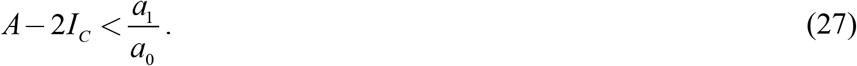

In this regime, perturbations due to a driving signal (e.g., random noise) around the fixed point of zero will lead to activity that returns to zero, with a rate constant given by the eigenvalue that is closest to zero. As a result, this regime will not have by prolonged periods of turning in one direction.

In the unstable regime (when the inequality in eq. (27) is reversed), behavior contrasts markedly. Perturbations around the fixed point will lead to activity that diverges until it limited by the sigmoid activation function of the full model, or driven back to zero by another perturbation. Thus, this regime will be characterized by prolonged periods of turning in one direction, or the other. The duration of these periods will be influenced by the extent of instability of the fixed point: with greater instability, the activity variables will be driven more strongly towards the limits of the sigmoid, and perturbations that suffice to drive them back to zero will be increasingly rare. As increasing *k*_*C*_ leads to greater instability, this behavior will be accentuated as *k*_*C*_ increases (i.e., as *I*_*C*_ becomes more negative). As shown by the simulations in the body and below, only slight instability is required to account for the level of turning observed experimentally.

#### Model regimes

Figure S3 illustrates how the locomotion trajectories depend on *A* and *I*_*C*_ for a range of values around the parameters chosen in the main text and that straddle the values at which the fixed points change their behavior. For *A* = 0.7 (top row), the fixed point of the antisymmetric component is stable for *I*_*C*_ ≥ −0.005 (first four panels with baseline and offset parameters, and fifth panel with baseline parameters). Under these circumstances, the trajectories are straighter than experimentally observed. For *A* = 0.9 (bottom row), the antisymmetric fixed point is unstable, and becomes progressively more so as *I*_*C*_ becomes more negative (from left to right). These trajectories have tight corkscrew components, which are not observed experimentally. For *A* = 0.8 (middle row), for values of *I*_*C*_ in the range −0.015 to −0.035, the models produce trajectories with modest curvature, consistent with experimental data. These parameters yield a fixed point that is slightly unstable. Similar trajectories are also seen in the rightmost panels of the top row, where also, the fixed point of the antisymmetric component is also slightly unstable. However, the underlying dynamics are quite different, because the fixed point of the symmetric component is stable. Consequently, there are no sustained periods in which the stop unit is active vs. inactive – so in this regime, the model fails to predict the experimentally-observed stops in the trajectory.

#### Role of the instability generated by contralateral inhibition

The analysis above shows that in the absence of contralateral inhibition, instability generated by stop inhibition or self-excitation cannot substitute as an explanation for trajectory curvature. Without contralateral inhibition (i.e., when *k*_*C*_ = *I*_*C*_ = 0), instability can be generated when *k*_*E*_ > 0 (corresponding to *A* > 0.8) or when *k*_*S*_ is sufficiently positive (*I*_*S*_ sufficiently negative), as shown by eqs. (23) and (25). However, this instability will not generate prolonged periods of turning in one direction, as the instability has its origin in the symmetric subsystem. Specifically, dynamics will fluctuate between periods in which there are extremes of activity in the symmetric subsystem (stop unit quiescent and other units active, or occasionally stop unit active and other units quiescent), but, since *I*_*C*_ = 0, there won’t be any tendency for bilaterally-paired units to be out of phase. Consequently, the swings of activity in the stop unit will interrupt periods of large forward velocity and periods of turning, but those periods of turning will have directions that fluctuate around zero, rather than maintained in one or the other direction.

Note also that a self-excitation value of zero (*k*_*E*_ = 0, or *A* = 0.8) positions the network’s behavior to be maximally sensitive to the level of contralateral inhibition. At this neutral point, the stability of the antisymmetric subsystem can be pushed into instability by the smallest changes in *I*_*C*_.

#### Model variants: one unit or multiple units on each side

The behaviors described above apply to model variants that have only one unit on each side, or more than two units, provided that they have the same connectivity pattern as the original model. The analysis is entirely parallel to the analysis detailed above, and we summarize it here.

We consider a model with *N* units on each side, each reciprocally connected to a single stop unit with inhibitory strength *k*_*S*_, and each reciprocally connected to all units on the opposite side with inhibitory strength *k*_*C*_. As in the original model, we postulate that summed signals from bilaterally-symmetric units control forward velocity, and differences control angular velocity.

The linearized model retains the form of eq. (6), and the connectivity pattern is embodied in a (2*N* +1)×(2*N* +1) matrix *L* given by

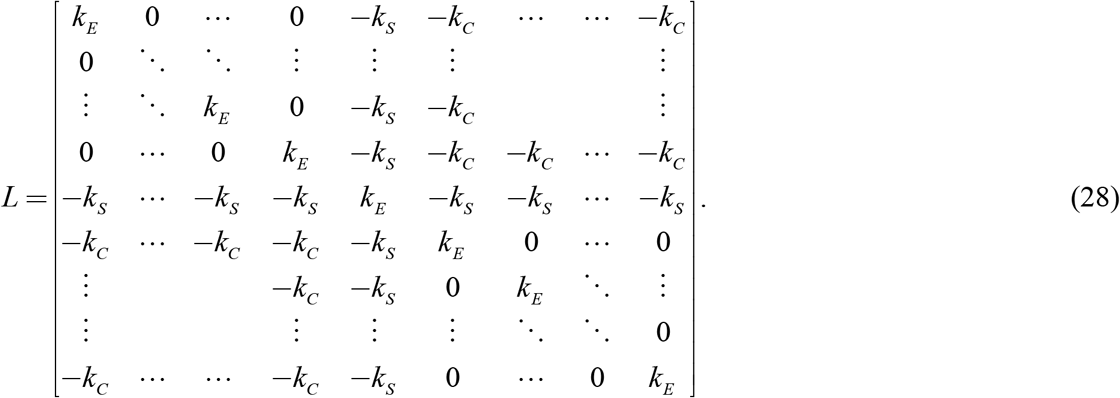

A rotation analogous to eq.(12) splits the system into a symmetric subsystem of dimension *N* +1 and an antisymmetric subsystem of dimension *N* (analogous to eqs. (17)), with

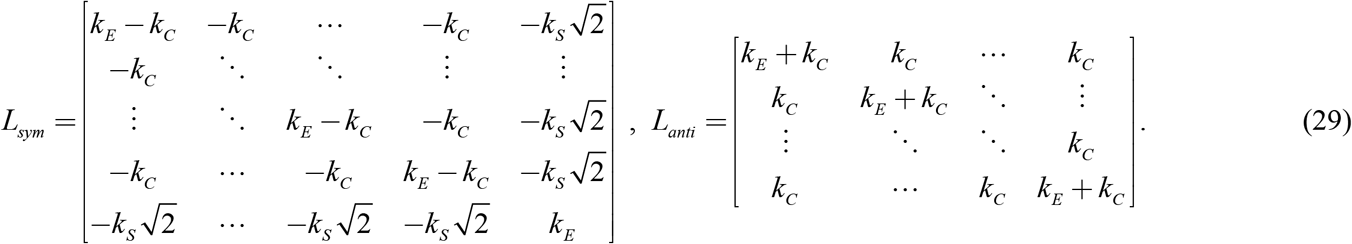

Via methods analogous to those used above, the symmetric subsystem is found to have *N* −1 eigenvalues equal to *k*_*E*_, and two other eigenvalues given by

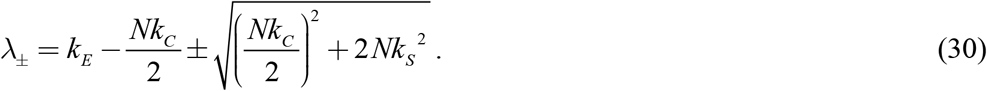

The antisymmetric subsystem has *N* −1 eigenvalues equal to *k*_*E*_ and one eigenvalue given by

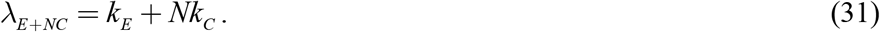

With *N* = 2, eq. (30) corresponds to eq. (23) and eq. (31) corresponds to eq. (26).

Thus, the stability behavior of a model with *N* units on each side is identical to that of the original model. In particular, *k*_*C*_ but not *k*_*S*_ contributes to instability in the antisymmetric subsystem. In detail, a model with *N* units on each side effectively multiplies the contralateral inhibitory strength *k*_*C*_ by *N*, multiplies the stop strength *k*_*S*_ by 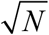, and creates *N* −1 eigenvalues equal to *k*_*E*_ in both subsystems.

#### Alternative connectivity, ipsilateral only

Here we analyze a model in which contralateral inhibition is replaced by ipsilateral connectivity (Figure S3 of the main text). In this alternative model, the only coupling between the two sides is through the stop neuron. As this model has the same symmetry as the ipsilateral-connectivity model, its analysis is closely parallel.

In the ipsilateral model, the connectivity matrix *L*_*ipsi*_ is given by

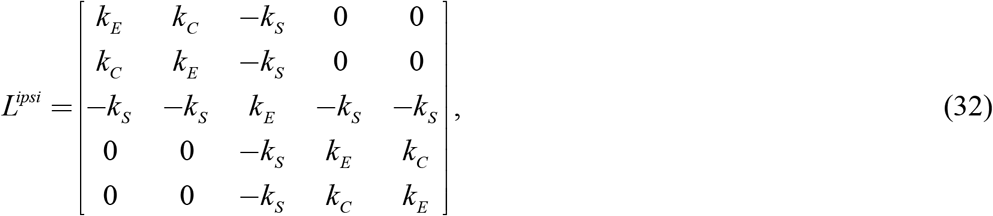

where the ipsilateral excitation strength (for ease of comparison with the original model) is denoted by *k*_*C*_ (positive for excitatory, negative for inhibitory; the correspondence to eq. 8 in the main text is 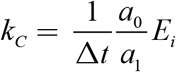). As in the analysis of the original model, the eigenvalues and eigenvectors can be determined by exploiting the model symmetery, and rotating *L*_*ipsi*_ into block-diagonal form.

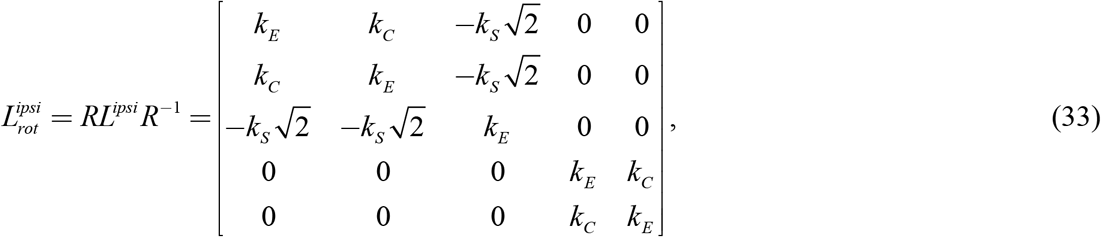

#### Symmetric component

The symmetric component,

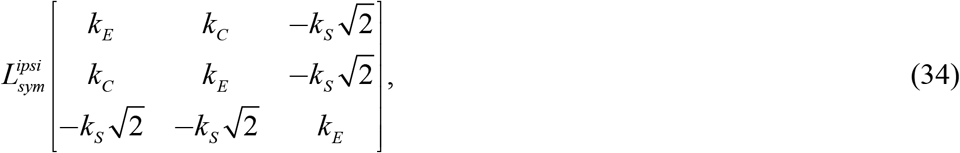

has an eigenvector [1 −1 0]′ with eigenvalue *λ*_*E*−*C*_ = *k*_*E*_ − *k*_*C*_, and two others, which must be orthogonal, of the form 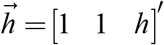. To determine *h*, note that if 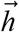 is an eigenvector, then 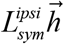 must be proportional to 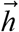, so the third element of

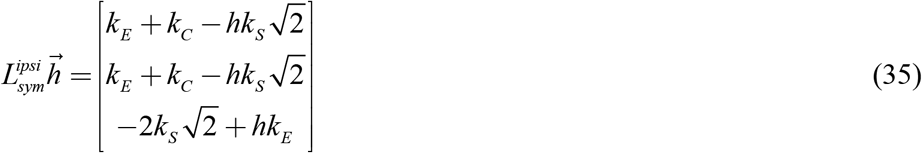

must be *h* times as large as each of the first two elements. This requires

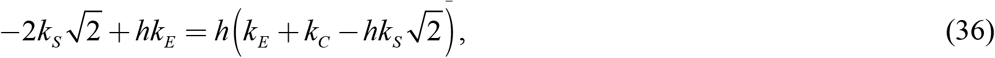

a quadratic equation whose roots are given by

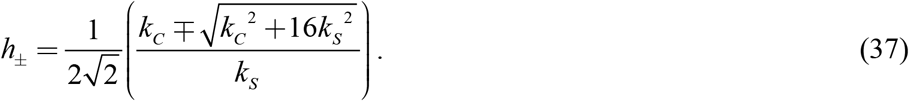

The eigenvalues are

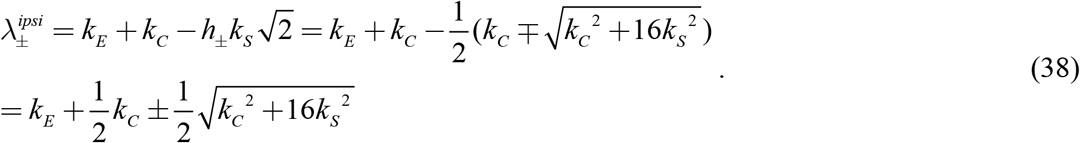

#### Antisymmetric component

The antisymmetric component,

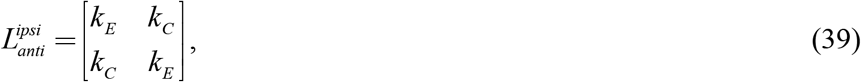

has eigenvectors [1 1]′ and [1−1]′ with eigenvalues

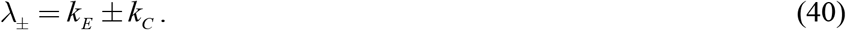

#### Comparison

Differences in the behaviors of the ipsilateral-connectivity model and the original model (contralateral inhibition) demonstrated by simulations in the main text (Figure S3) are accompanied by changes in how the stability characteristics of the fixed point depends on connectivity parameters (see Table below). We focus on their behaviors when *k*_*E*_ is near zero (the range considered in the text) and a nonzero stop-inhibition *k*_*S*_. *k*_*C*_ > 0 corresponds to contralateral inhibition in the original model, or ipsilateral excitation in the alternative model.

In the original model, for *k*_*E*_ near zero and *k*_*C*_ > 0, both subsystems have a neutrally stable eigenvalue and an unstable one, and the symmetric subsystem also has a stable eigenvalue. Increasing *k*_*C*_ increases the instability in the antisymmetric system (increasing the duration of bouts of turning and typical angular velocities) but decreases the instability in the symmetric system (decreasing forward velocities), corresponding to the behavior seen in Figure 3E.

In the models with ipsilateral connectivity, this opposite influence of *k*_*C*_ on the stability characteristics of the two subsystems does not occur. Both subsystems have a stable and an unstable eigenvalue, and the symmetric subsystem has a smaller-magnitude third eigenvalue whose characteristics depend on the sign of *k*_*C*_. Increasing the magnitude of *k*_*C*_ merely moves the most positive and most negative eigenvectors further from zero. Correspondingly, the effects on forward and angular velocity (Figure S3C and S3F) are in the same direction, rather than opposite.

